# Eye-head coordination during goal-directed orienting in mice

**DOI:** 10.64898/2026.03.30.715285

**Authors:** Brandie Morris Verdone, Hui Ho Vanessa Chang, Dale C. Roberts, Kathleen E. Cullen

## Abstract

In afoveate species such as mice, it is accepted that gaze is typically redirected by head movements with a saccade-and-fixate strategy, while the eyes primarily stabilize vision within a limited oculomotor range. This view suggests that the accompanying eye movements are primarily reflexive, driven by mechanisms like the vestibulo-ocular reflex (VOR). However, emerging evidence challenges this assumption, suggesting that eye movements during active head motion may not be purely reflex-driven. Here, we directly test whether eye movements in mice are actively coordinated as part of voluntary gaze redirection rather than being reflexive. By systematically monitoring head and pupil positions during goal-directed orienting in a cohort of male mice, we find that mice generated active saccadic eye movements whose onsets are tightly linked to head movements. Furthermore, these saccadic eye movements occur at markedly shorter latencies than reflexive quick-phase eye movements evoked by comparable passive head rotations. Importantly, the interplay between coordinated eye and head movements during voluntary orienting resemble the predictable, stereotyped gaze patterns seen in foveate animals, such as primates. Our results suggest that mice possess an evolutionarily conserved mechanism for gaze redirection, integrating voluntary eye-head coordination similar to that of foveate vertebrates. These findings reframe the prevailing view by demonstrating an actively coordinated eye-head component to gaze redirection under goal-directed conditions in mice, complementing established reflexive mechanisms.

## INTRODUCTION

Humans and other foveate species, such as non-human primates, commonly make voluntary coordinated eye and head movements to redirect their gaze from one part of the visual field to another^1–7^. These gaze shifts are characterized by saccades (rapid eye movements that redirect the point of visual focus from one target to another) made in tandem with corresponding movements of the head and, at times, even the entire body^4–8^. Because gaze is determined by the sum of eye position relative to the head and head position relative to space, accurate fixation requires the precise coordination of head and eye movements. The combination of these orienting movements with intermittent periods of gaze stabilization comprises a ubiquitous pattern of voluntary movement that is essential for navigating and interacting with the environment^6, 9, 10^; this has been termed a “saccade and fixate” strategy^11^.

In contrast to foveates, most vertebrates lack a fovea—the retinal region of highest visual acuity, composed exclusively of cone photoreceptors—and are thus termed *afoveates*. When exploring the environment, afoveates also use a “saccade and fixate” strategy^11^. A prevailing view has been that the eye movements made by afoveates when implementing this strategy serve to maintain stable gaze rather than actively redirect their gaze to target specific objects^6, 12^. In this view, two essential eye movement reflexes that evolved in early vertebrates—the vestibulo-ocular reflex (VOR) and optokinetic reflex (OKR)—work synergistically to keep the visual world fixed on the retina by moving the eye counter to head motion. Notably, in response to prolonged head motion, these reflexes initially produce stabilizing eye movements that counteract head motion (slow-phases), followed by rapid eye movements in the direction of head motion that recenter the eye (quick-phases). As these quick-phases tend to move the average position of the eye in the direction of passive head movement (i.e. beating field), it has been proposed that they make a functional contribution to the orienting response^13, 14^.

Mice are an afoveate vertebrate species widely used as a model in vision research. As afoveates, mice are generally considered to use a “saccade and fixate” strategy^15, 16^, where intentional scanning head movements induce reflexive eye movements to ensure stable gaze during exploration^17–20^. This proposal aligns with observations that mice, like most vertebrate species, generate robust slow and quick-phase VOR and OKR eye movements^21, 22^. While this viewpoint has informed the interpretation of recent studies on gaze strategy in the mouse^16, 18, 19, 23^, studies have also challenged the notion that the eye movements made during scanning head movements are driven exclusively by reflex pathways. For example, the superior colliculus (SC), which is essential for active gaze control^24, 25^, induces eye and head movements in monkeys (foveates) and cats (afoveates) when stimulated^8, 26–30^. Furthermore, recent evidence indicates that stimulating the SC in mice also results in significant eye^23^ as well as head^31^ movements similar to those observed in cats and monkeys. Indeed, when mice were observed during free movement, conjugate, saccadic horizontal eye movements were seen alongside attempted head movements^18^. Thus, it stands to reason that there may exist a common neural strategy for eye-head coordination that is conserved across different species, in foveates and afoveates alike.

Accordingly, here we directly investigated if eye movements in mice comprise a coordinated component of voluntary gaze redirection. We took advantage of a well-defined training paradigm in which head and pupil position were tracked while mice made active, goal-directed orienting head movements. We found that mice, despite being afoveate, generated active saccadic eye movements in coordination with these orienting head movements. Notably, we established that these active, saccadic eye movements are directed in the same direction as the active head movement and occur at markedly shorter latencies than the reflexive, quick-phase eye movements evoked by comparable passive head rotations. The onset of these saccadic movements was tightly linked to that of head motion, with a significant proportion preceding head motion onset. Furthermore, the interplay between eye and head movements during active orientation in mice mirrored the patterns seen in primates, maintaining certain predictable relationships; for example, eye movements initiated with the eye more contralateral in the orbit were accompanied by smaller head movements. Taken together, our findings suggest the presence of conserved features in the generation of voluntary eye-head gaze shifts across diverse species, supporting the notion of a shared evolutionary mechanism for gaze redirection.

## RESULTS

### Establishing an active orienting paradigm in mice

To understand the coordination of eye and head motion during active orienting movements, mice were trained to perform a reward-based behavioral assessment similar to a task used in prior studies of cat gaze shifts^32–34^. Nine male mice were body-restrained in a tube and attached via their headposts to a custom interface that allowed them to move their heads freely in the horizontal (yaw) plane, similar to devices used in non-human primate studies^4, 7, 35^. A high precision potentiometer was used to measure angular head position **(Figure 1A)**, and a custom lightweight head-mounted camera system was positioned in front of the eye to track the pupil using video oculography (VOG) software **(Figure 1A, C)**. To evoke active orienting head movements of different sizes, the waterspouts were positioned at ±10° from center for small, ±25° for intermediate, and ±40° for large head movements **(Figure 1B)**. Upon the onset of a randomized tone, the mouse moved its head to a spout to receive a water reward. When the tone repeated, the mouse was then required to redirect its head to the opposite spout for a reward, as no reward would be given for incomplete movement (e.g., pausing in-between spouts at the onset of a tone or failure to move their head to the opposing spout). This reward paradigm, combined with head and eye position tracking, allowed for the assessment of the natural coupling between eye and head movements as mice engaged in goal-directed behavior, as would more typically occur during orienting tasks in natural settings. In addition to this active orienting paradigm, we also manually applied an external torque to each mouse’s head that passively rotated it between the same small, intermediate, and large angular distances in a separate paradigm **(Figure 1A)**. Together, comparison across these paradigms allowed for a direct assessment of differences in eye and head coordination during active orienting versus passive head movements.

**Figure 1:**
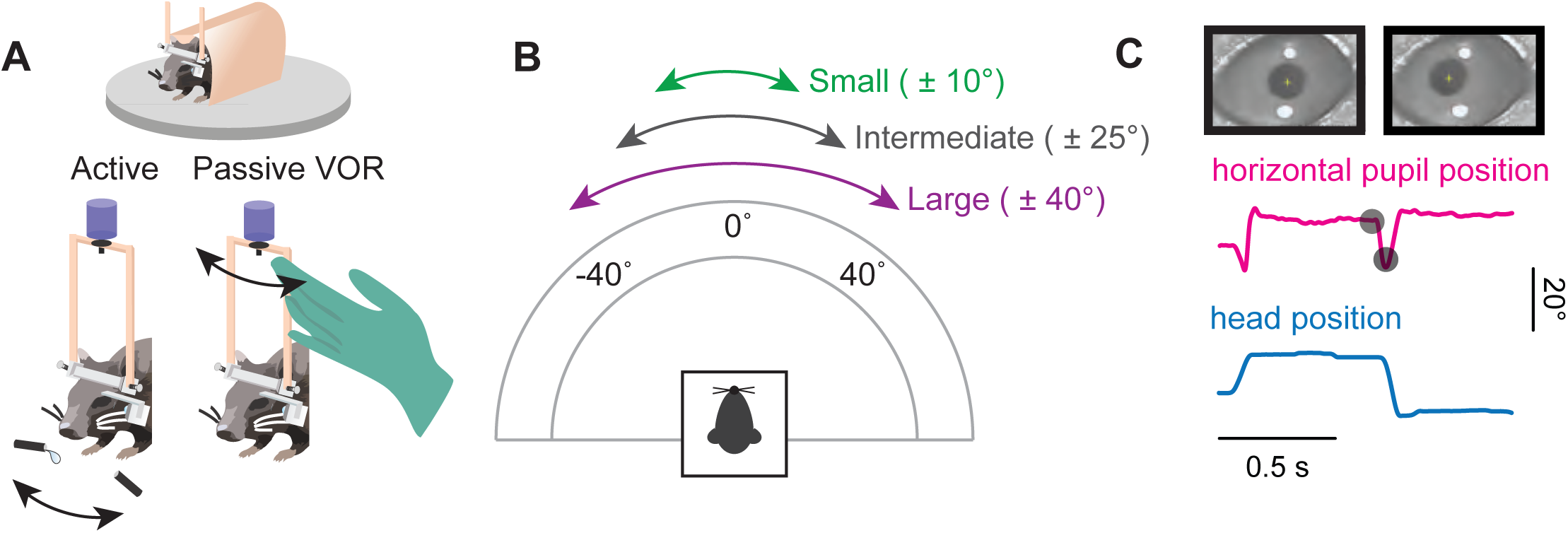
Experimental Design. **A**. After undergoing headpost and camera mount implantation surgery (see Methods), the mouse was body-restricted in a tube and its headpost was connected to a potentiometer to measure head position. A small camera was attached to the camera mount to capture the mouse’s pupil. In the active paradigm, the mouse voluntarily oriented along the horizontal axis between two waterspouts for reward at the sound of a randomized tone. For the passive VOR paradigm, the experimenter rotated the mouse’s head. **B.** Schematic of angular distances along the horizontal axis. For active, the mouse oriented its head between waterspouts for ± 10°, ± 25°, and ± 40° for small, intermediate, and large angular distances. For passive VOR, the experimenter rotated the mouse’s head between ± 10°, ± 25°, and ± 40°. **C.** Representative traces of head position (blue) and horizontal pupil position (magenta) during a segment of the active orienting task. Black dots correspond to the pupil positions in the still frames. Vertical scale bar: 20° Horizontal scale bar: 0.5 seconds. Also see Supplementary Figure 1.

### Characteristics of eye and head coordination during active orienting

Following successful training (see **Methods, Supplementary Figure 1**), mice performed the active paradigm with simultaneous head and eye position tracking for assessment of their dynamics and respective contributions to gaze. We determined the onset of head, eye, and overall gaze movement by computing the time at which the respective velocities reached a threshold of 25 degrees per second in the direction (temporal or nasal) of the orienting movement.

If, during active exploration, mice generated orienting head movements that elicited compensatory eye movements to stabilize gaze, then the onset of head motion would be expected to precede gaze onset (“Head-Initiated”). This occurs because the compensatory slow phase of the vestibulo-ocular reflex (VOR) initially drives the eyes in the opposite direction to the orienting head movement. As a result, gaze redirection is delayed until the quick phase of the VOR generates a rapid eye movement in the same direction as the head. Consistent with this expectation, such “head-initiated” movements were observed across all tested amplitudes, where the eye rapidly followed the head once gaze redirection began (**Figure 2A–C**).

**Figure 2:**
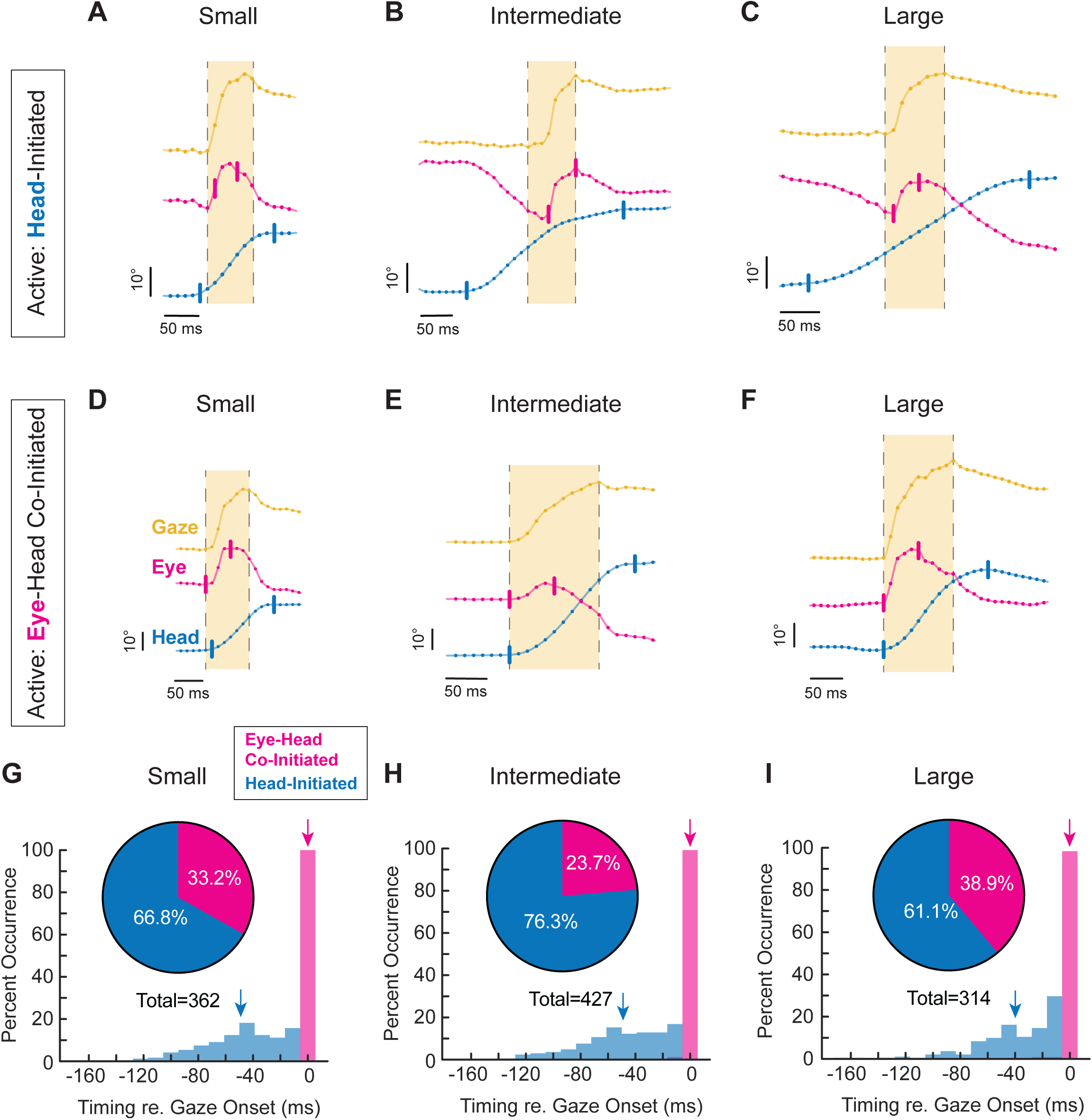
Position, distribution, and timing of head-initiated vs. eye-head co-initiated orienting. **A-C.** Head-initiated orienting for small, intermediate, and large orienting movements, respectively, depicted as position. **D-F.** Eye-head co-initiated orienting for small, intermediate, and large orienting movements, respectively, depicted as position. Yellow = gaze. Magenta = eye. Blue = head. Vertical tick marks indicate onset and offset of head or eye, respectively. Onset and offset cutoffs of 25 °/s. Light yellow window indicates gaze duration. Vertical scale bar: 10°. Horizontal scale bar: 50 milliseconds. Dots indicate 90 Hz sampling, with positional traces up-sampled to 1kHz for visualization. **G-I.** Distribution (insets) and timing with respect to gaze onset of small, intermediate, and large head-initiated and eye-head co-initiated orienting movements. Magenta = eye-head co-initiated, Blue = head-initiated. Arrows indicate mean of respective signal. **G.** Small inset: 66.8% head-initiated versus 33.2% eye-head co-initiated. Mean latency head: -49.2±29.9 milliseconds. Mean latency eye: 0.0 ± 0.0 milliseconds. Head-initiated v. eye-head co-initiated p < 0.0001. N = 362 segments across 9 mice. **H.** Intermediate inset: 76.3% head-initiated versus 23.7% eye-head co-initiated. Mean latency head: -49.3± 30.8 milliseconds. Mean latency eye: -0.11 ± 1.1 milliseconds. Head-initiated v. eye-head co-initiated p < 0.0001. N = 427 segments across 9 mice. **I.** Large inset: 61.1% head-initiated versus 38.9% eye-head co-initiated. Mean latency head: -39.3 ± 29.5 milliseconds. Mean latency eye: -0.27 ± 2.2 milliseconds. Head-initiated v. eye-head co-initiated p < 0.0001. N = 314 segments across 9 mice. Histogram statistical comparisons performed using the Wilcoxon rank-sum test. Latencies shown as mean ± SD. Histogram bin size = 11.1 milliseconds, corresponding to a 90 Hz sampling rate, or 1 video frame. Also see Supplementary Figure 2, Supplementary Figure 3, Video S1, and Video S2. Bimodal distributions were observed for small, intermediate, and large head-initiated gaze redirections (blue, Hartigan’s dip test, p<0.0005).

Unexpectedly, however, we also commonly observed orienting movements in which gaze onset either preceded or coincided with head motion (“Eye–Head Co-Initiated”; see Methods). In these cases, a distinct rapid eye movement in the direction of the orienting movement occurred just before or nearly simultaneously with head motion onset. Because such movements reach onset threshold before head motion, they cannot be driven by stabilizing reflexes, which depend on vestibular input to counteract head movement. This close temporal coupling between eye and head onset therefore suggests a coordinated motor strategy, in which gaze onset was initiated by a rapid eye movement in the same direction as the head (**Figure 2D–F**).

Overall, our analysis revealed that while active orienting was generally initiated by head movements, eye-head co-initiated gaze redirections were likewise observed across all angular distances **(Figure 2A-C)**. Across head-initiated orienting movements, head motion onset preceded gaze onset by an average of 49.2 ± 29.9 milliseconds for small **(Figure 2G, blue arrow)**, 49.3±30.8 milliseconds for intermediate **(Figure 2H, blue arrow)**, and 39.3±29.5 milliseconds for large orienting movements **(Figure 2I, blue arrow)**. This pattern is consistent with an initial recruitment of the compensatory slow-phase of VOR, during which the eye initially moves opposite the head, preventing gaze from reaching threshold. In contrast, for the subset of orienting movements co-initiated by eye and head, eye and gaze onset were tightly linked with an average interval of 0.0±0.0 milliseconds, 0.1±1.1 milliseconds, and 0.3±2.2 milliseconds for small, intermediate, and large orienting movements, respectively **(Figure 2G-I, magenta arrows)**. Although there was a propensity for head motion onset to precede gaze, eye-head co-initiated gaze redirections were routinely observed across movement amplitude (33.2%, 23.7% and 38.9%, respectively; **Figure 2G-I, insets)**. In addition, the head-initiated distributions in Figure 2G-I were significantly bimodal across small, intermediate, and large amplitudes (Hartigan’s dip test, p < 0.0005). This bimodality suggests the presence of at least two temporal strategies in the timing of head-initiated gaze redirections —one characterized by a shorter head lead and another by a longer head lead—consistent with differential VOR engagement near gaze onset in head-initiated trials.

We next tested whether our findings might be influenced by mice attempting to move their heads via enbloc movements of their head-neck ensemble rather than movements of the head itself about the cervical column. To address this possibility, we conducted an experiment where we further restricted the lumbar region of the mouse’s body during our active behavior paradigm. This comparative condition (see Method Details: Active and Passive Behavioral Paradigms) ensured that head movement occurred with reduced rotation about the back of the body. Interestingly, in this more restricted condition, we observed an even larger percentage of eye-head co-initiated gaze redirections for small and intermediate gaze shifts (53.7% and 39.3% respectively) (**Supplementary Figure 2**). The percent occurrence of eye-head co-initiated gaze redirections during large orienting movements was comparable to the original testing condition (29.6%). Together, this suggests that reducing lumbar contribution to attempted head movements results in increased or comparable gaze redirections co-initiated by eye-head movement.

As our setup was specifically designed to restrict orienting movements to the horizontal (yaw) axis, we wanted to confirm that this restriction did not affect the relative timing of rapid eye movements to orienting head movements. Thus, we conducted an additional experiment in which head motion was tracked using a miniaturized lightweight inertial measurement unit (IMU) with the head restriction removed (‘Head-Free’, see Methods: Active and Passive Behavioral Paradigms). We first confirmed that the addition of an IMU when performing the initial, yaw-restricted, paradigm did not alter the distribution of head-initiated versus eye-head co-initiated gaze shifts (**Figure 2G; Supplementary Figure 3A**; χ²=.0245, dof=1, p=0.876). We then assessed performance in the head-free condition and again found that the relative proportion of head-initiated versus eye-head co-initiated orienting movements was consistent with that observed in the yaw-restricted paradigm. Notably, 31.2% of unrestricted orienting movements were eye-head co-initiated (**Supplementary Figure 3B)**, comparable to the 34.6% observed during yaw-restricted orienting (**Supplementary Figure 3A;** χ²=0.169, dof=1, p=0.681). Together, these results demonstrate that active orienting head movements are closely synchronized with the onset of rapid eye movements in the same direction, facilitating gaze redirection. Interestingly, the mean eye velocity was higher in the head-free condition than in the yaw-restricted condition (**Supplementary Figure 3D**), despite comparable mean head velocities across conditions (**Supplementary Figure 3C**), suggesting that faster eye movements are employed when the head is unconstrained.

### Eye and head contributions to active orienting

To better understand the head-eye gaze coordination outlined in **Figure 2**, we next visualized the head, eye, and gaze redirection generated during active orienting in terms of velocity rather than position. **Figure 3** highlights the trajectories of head (blue), eye (magenta), and gaze (yellow) over a series of orienting movements ranging from small to large made by a mouse during our paradigm. As seen across all mice in the study **(Supplementary Figure 4)**, relatively stereotyped head, eye, and gaze trajectories were observed when aligned to gaze onset **(Figure 3A, Supplementary Figure 4A)**. Overall, we found that specific head-eye dynamics emerged depending on whether an active orienting movement was head-initiated or eye-head co-initiated. In scenarios where gaze redirection was co-initiated by the eye and head, both the eye and head moved ipsilaterally and an initial VOR engagement was absent **(Figures 3B, D, Supplementary Figure 4D).** Meanwhile, VOR engagement was observed in conditions where a rapid eye redirection occurred following an initiating head movement **(Figures 3C, E, Supplementary Figure 4E)**. All active orienting movements terminated with opposing head and eye movements, indicative of reflexive compensation to stabilize gaze at the end of a redirection. These patterns of eye-head movements were consistent with the latency estimations shown in **Figure 2 G-I**; an increased head lead time suggests that the eye initially moves in the opposite direction as the head, while tight eye-gaze latency suggests a short period between eye and head movement in the same direction. Both strategies (head-initiated and eye-head co-initiated) were observed across all mice (**Supplementary Figure 4)**, indicating that the presence of eye-head co-initiation was not driven by a small subset of subjects. Subtle between-animal differences in eye, head, and gaze trajectories were observed across mice (**Supplementary Figure 4A-E**). In addition, the relative prevalence of these strategies varied significantly across animals (**Supplementary Figure 4F**; χ² = 84.09, dof = 8, p < 0.0001). Similar inter-individual differences in head-eye gaze coordination have been reported in monkeys performing a comparable orienting task^36^.

**Figure 3:**
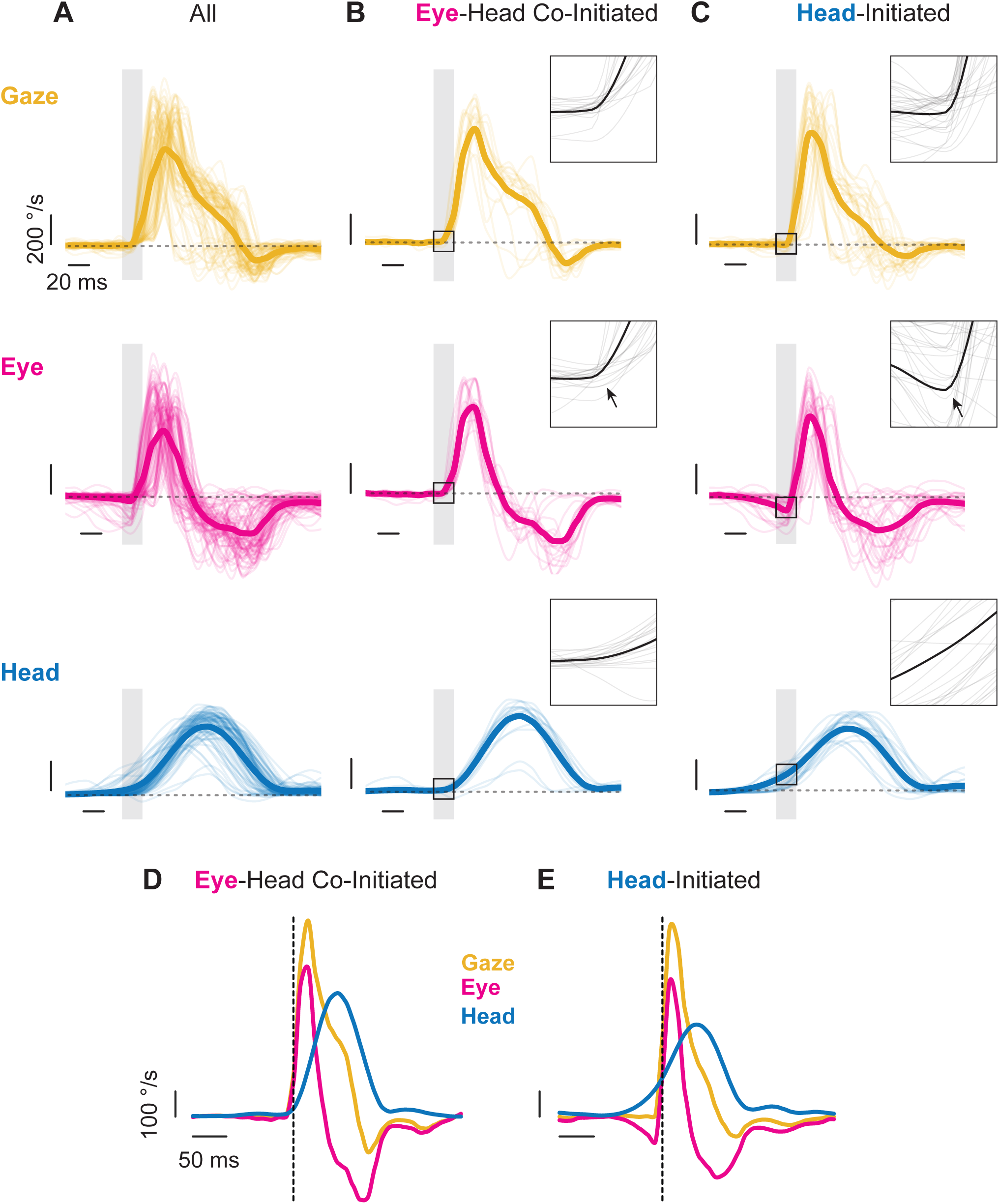
Head, eye, and gaze redirection velocities. Orientations as shown in velocity traces of all **(A)**, eye-head co-initiated **(B)**, and head-initiated **(C)** orienting movements in a single direction from a representative mouse. Eye and head onsets aligned to gaze onset. Yellow = gaze. Magenta = eye. Blue = head. Individual traces shown as thin lines. Means of respective signal as thick line. Grey windows and black box highlight a 20-millisecond region of interest around gaze onset, highlighting VOR engagement for head-initiated gaze redirection and the absence thereof for eye-head co-initiated gaze redirection (arrows). **A-C.** Vertical scale bar: 200 °/s. Horizontal scale bar: 20 milliseconds. Insets are enlargements of the 20-millisecond region of interest highlighted by the black box. N = 45 segments. Dashed grey line at 0 °/s. Mean of eye-head co-initiated **(D)** and head-initiated **(E)** traces shown. **D, E.** Vertical scale bar: 100 °/s. Horizontal scale bar: 50 milliseconds. Velocity traces up-sampled to 1 kHz for visualization. Vertical dashed line indicates gaze onset. Also see Supplementary Figure 4.

Next, we assessed the relative contributions of eye and head to gaze for these active orienting movements. **Figure 4A-D** illustrates the relationships between head and eye amplitude (range of movement) with respect to the amplitude of the overall change in gaze. Movements in both the nasal and temporal directions showed a linear relationship that increased from small to intermediate to large active orientations. Both temporally and nasally directed head movements showed a statistically significant contribution to gaze as measured in amplitude **(Figure 4A, B)** (Linear Regression, Head Temporal: R^2^=0.63, p<0.0001; Head Nasal: R^2^=0.59, p<0.0001). Eye amplitude in both the temporal and nasal directions increased with gaze amplitude in a manner that was well described by an exponential function **(Figure 4C, D)** (Eye Temporal: R^2^=0.38, τ=21.2°; Eye Nasal: R^2^=0.37, τ=14.5°). Further, the average horizontal eye amplitude during active orientation was 10.6 ± 4.8°, 10-fold larger than the corresponding vertical eye amplitude (0.9 ± 0.9°). Despite the contribution of the eye being statistically significant, the amplitude of head movements had a larger contribution to the overall change in gaze, in line with primate studies^4, 7^. When stratified by gaze redirection type (head-initiated versus eye-head co-initiated, **Supplementary Figure 5 A-D**), modest but significant differences in main sequence relationships emerged. Specifically, in the temporal direction, head movement amplitude scaled more steeply with gaze amplitude during eye-head co-initiated orienting movements (slope = 1.17, R² = 0.68) than during head-initiated movements (slope = 0.98, R² = 0.56; p = 0.01). A similar difference was observed for eye movement amplitudes, with eye-head co-initiated movements characterized by a smaller exponential constant (τ = 22.0°, R² = 0.57) than head-initiated movements (τ = 25.0°, R² = 0.37; p < 0.0001).

**Figure 4:**
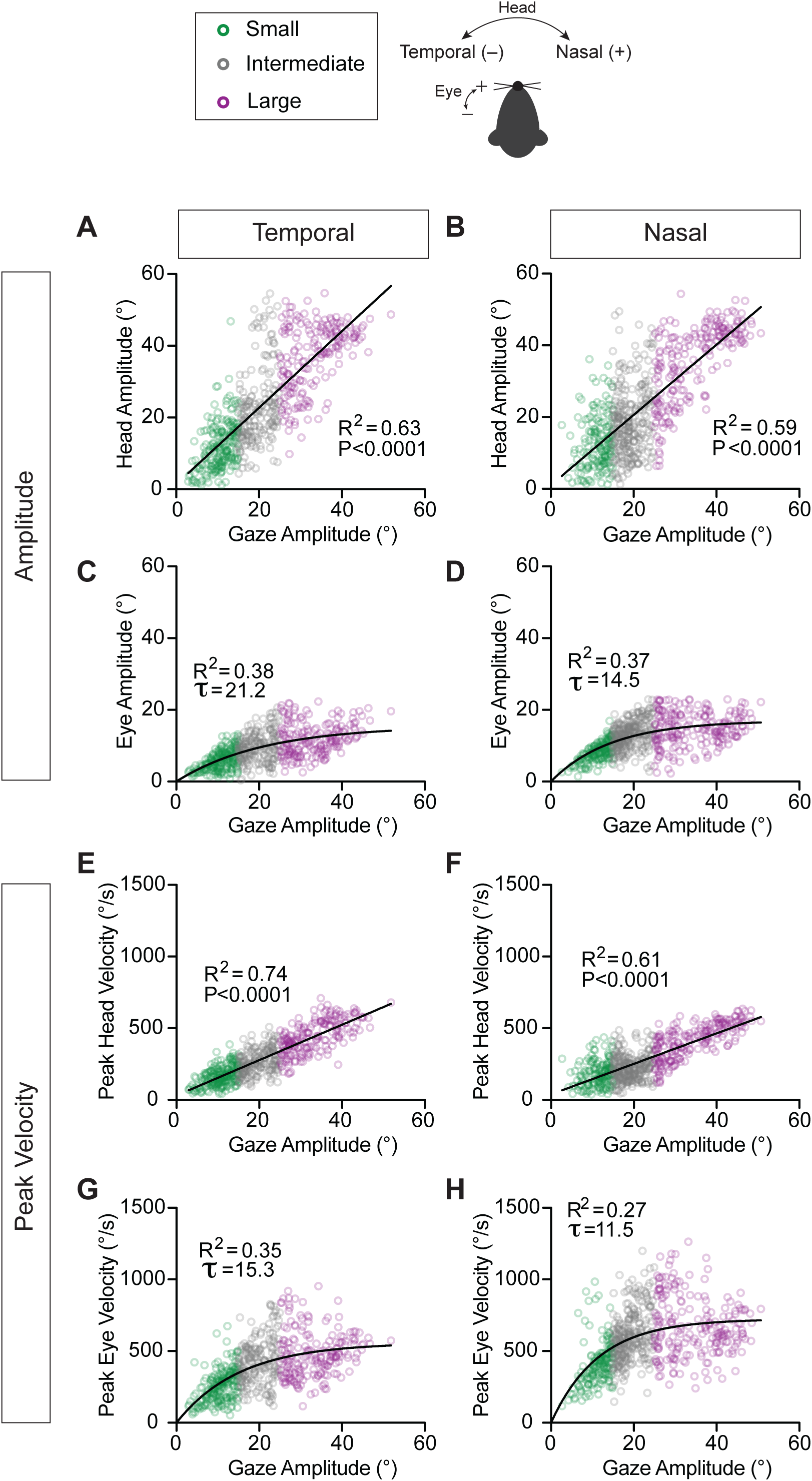
Head and eye amplitudes and peak velocities with respect to gaze amplitude. A, **B.** Head movement amplitude with respect to gaze amplitude for temporally **(A)** and nasally **(B)** directed movements. Linear Regression, Head Temporal: R^2^ = 0.63, p < 0.0001, slope=1.07 (95% CI: [1.0, 1.13]), N = 581 segments across 9 mice; Head Nasal: R^2^ = 0.59, p < 0.0001, slope=0.98 (95% CI: [0.92, 1.04]), N = 640 segments across 9 mice. **C, D.** Eye movement amplitude with respect to gaze amplitude for temporally **(C)** and nasally (**D)** directed movements. Eye Temporal: R^2^ = 0.38, τ = 21.2° (95% CI: [17.2, 26.7]), N = 581 segments across 9 mice; Eye Nasal: R^2^ = 0.37, τ = 14.5° (95% CI: [12.7, 16.6]), N = 638 segments across 9 mice. **E, F.** Peak velocity of head movement with respect to gaze amplitude for temporally **(E)** and nasally **(F)** directed movements. Linear Regression, Head Temporal: R^2^ = 0.74, p < 0.0001, slope =12.3 (95% CI: [11.8, 12.9]), N = 581 segments across 9 mice; Head Nasal: R^2^ = 0.61, p < 0.0001, slope =10.6 (95% CI: [9.98, 11.3]), N = 640 segments across 9 mice. **G, H**. Peak velocity of eye movement with respect to gaze amplitude for temporally **(G)** and nasally (**H)** directed movements. Eye Temporal: R^2^ = 0.35, τ = 15.3° (95% CI: [12.9, 18.3]), N = 581 segments across 9 mice; Eye Nasal: R^2^ = 0.27, τ = 11.5° (95% CI: [10.0, 13.1]), N = 640 segments across 9 mice. Green = small, grey = intermediate, purple = large orienting movements. Schematic of temporal and nasal directions for head and eye shown.

To characterize eye and head movement kinematics during gaze redirection, we examined how peak eye and head velocities scaled with gaze amplitude **(Figure 4E-H)**. Indeed, we found that larger amplitude changes in gaze were accomplished via the generation of eye as well as head movements with increased velocities. Peak head velocities showed a linear increase with respect to gaze amplitude in both the temporal **(Figure 4E)** and nasal **(Figure 4F)** directions (Linear Regression, Head Temporal: R^2^=0.74, p<0.0001; Head Nasal: R^2^=0.61, p<0.0001). When stratified by gaze redirection type **(Supplementary Figure 5E)**, this relationship differed significantly for temporally directed movements: the slope for eye–head co-initiated orienting movements (slope=13.4, R^2^=0.79) was higher than that for head-initiated movements (slope=10.5, R^2^=0.66, p<0.0001). Peak eye velocities were again well described by an exponential function **(Figure 4G, H**) (Eye Temporal: R^2^=0.35, τ=15.3°; Eye Nasal: R^2^=0.27, τ=11.5°). Interestingly, the time constant (τ) of the nasally directed eye movements (**Figure 4H**, τ = 11.5°) was shorter than that of the temporally directed eye movements (**Figure 4G**, τ = 15.3°). This indicates that peak eye velocity rises more rapidly with gaze amplitude for nasal movements compared to temporal movements, consistent with prior reports of rapid eye movements made in the absence of head movement^37^, as well as head-unrestrained conditions^18^.

Consistent with this, eye-head co-initiated orienting movements exhibited a shorter time constant in the nasal direction (τ=11.3°) than head-initiated movements (τ=14.8°) (**Supplementary Figure 5H**). Overall, the velocities and amplitudes of both head and eye movements demonstrate predictable relationships with gaze during this active orienting behavior.

### Influence of initial eye position on head and eye contributions to gaze

Previous studies of gaze shift dynamics in humans and monkeys have emphasized the role that the starting position of the eye-in-orbit plays in head and eye coordination^4, 7, 38^. The eye’s contribution to gaze is greater when the initial eye position is contralateral to the direction of the intended gaze shift^4, 39–42^. We predicted that this pattern of head-eye coordination would also exist in mice and found that this was indeed the case. **Figure 5** shows the eye and head contributions to gaze (eye and head amplitudes) as a function of initial eye position. The head contribution to gaze increased as the initial eye position shifted towards the direction of the orienting movement—that is, the head contributed more when the eye was already positioned towards the intended direction **(Figure 5A, B, Supplementary Figure 6A, B)** (Linear Regression, Head Temporal—small R^2^=0.07, p=0.0002, intermediate: R^2^=0.27, p<0.0001, large: R^2^=0.43, p<0.0001; Head Nasal—small R^2^=0.15, p<0.0001; intermediate: R^2^=0.30, p<0.0001; large: R^2^=0.45, p<0.0001). Accordingly, the contribution of the eye to gaze increased when the initial eye position was directed opposite to the direction of the orienting movement **(Figure 5C, D, Supplementary Figure 6C, D)** (Linear Regression, Eye Temporal—small R^2^=0.08, p<0.0001, intermediate: R^2^=0.28, p<0.0001, large: R^2^=0.38, p<0.0001; Eye Nasal—small R^2^=0.05, p=0.0032; intermediate: R^2^=0.14, p<0.0001; large: R^2^=0.32, p<0.0001). This finding is consistent with work in freely moving mice showing that superior colliculus stimulation-evoked saccades are biased by initial eye position, such that saccades are more likely when initiated from positions opposite the intended endpoint^43^. Further, in visualizing the head and eye contributions to changes in gaze, it was clear that the relative relationships between small, intermediate, and large orienting movements were maintained, reminiscent of the relationship between head-eye amplitude and gaze depicted in **Figure 4 (Supplementary Figure 6, insets).**

**Figure 5:**
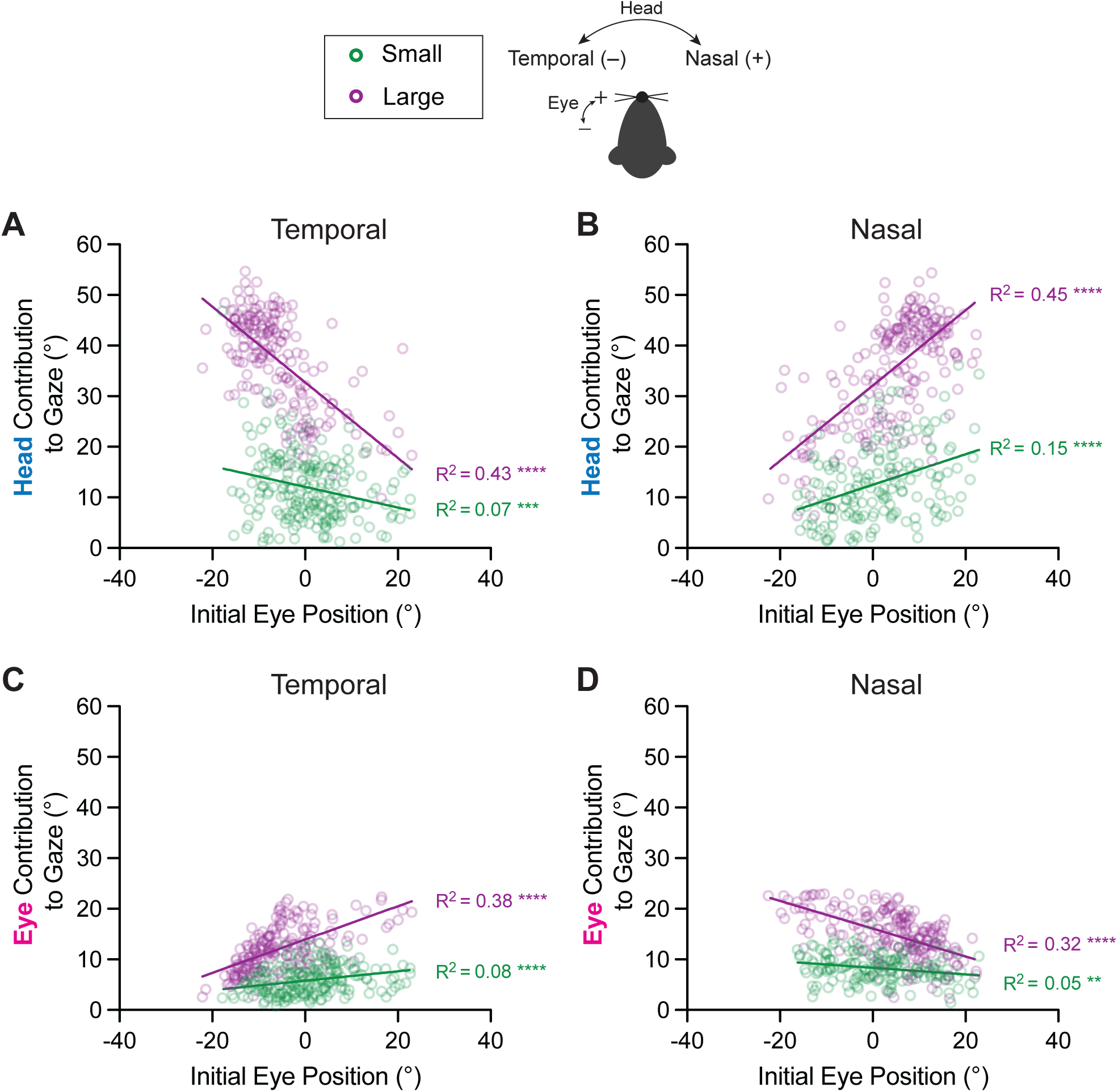
Influence of initial eye position on head and eye contributions to gaze. **A, B.** Contribution of head movement to gaze with respect to initial eye position for temporally **(A)** and nasally **(B)** directed movements. Linear Regression, Head Temporal—small R^2^ = 0.07, p = 0.0002, slope = -0.20 (95% CI: [-0.31, -0.10]), N = 205 segments across 9 mice; large: R^2^ = 0.43,, p < 0.0001, slope = -0.74 (95% CI: [-0.87, -0.62]), N = 190 segments across 9 mice. Line comparisons: F = 43.3, DFn = 1, DFd = 391, p < 0.0001; Head Nasal—small R^2^ = 0.15, p < 0.0001, slope = 0.30 (95% CI: [0.19, 0.41]), N = 164 segments across 9 mice; large: R^2^ = 0.45, p < 0.0001, slope = 0.74 (95% CI: [0.63, 0.86]), N = 209 segments across 9 mice. Line comparisons: F = 29.1, DFn = 1, DFd = 369, p < 0.0001. **C, D.** Contribution of eye movement to gaze with respect to initial eye position for temporally **(C)** and nasally **(D)** directed movements. Eye Temporal—small R^2^ = 0.08, p < 0.0001, slope = 0.09 (95% CI: 0.04, 0.14]),N = 205 segments across 9 mice; large: R^2^ = 0.38, p < 0.0001, slope = 0.33 (95% CI: [0.27, 0.39]),N = 190 segments across 9 mice. Line comparisons: F = 39.6, DFn = 1, DFd = 391, p < 0.0001; Eye Nasal—small R^2^ = 0.05, p = 0.0032, slope = -0.07 (95% CI: [-0.10, -0.02]), N = 164 segments across 9 mice; large: R^2^ = 0.32, p < 0.0001, slope = -0.27 (95% CI: [-0.33, -0.22]),N = 209 segments across 9 mice. Line comparisons: F = 32.48, DFn = 1, DFd = 369, p < 0.0001. Green = small, purple = large orienting movements. Schematic of temporal and nasal directions for head and eye shown. Also see Supplementary Figure 5.

### Active gaze redirection differs from passive gaze stabilization in mice

As reviewed above, a prevailing view suggests that mice make orienting head movements that induce reflexive eye movements to stabilize gaze. If so, rapid eye movements made during active head motion should mirror those made during passive head movement with comparable vestibular input. To test this, we directly compared head-eye relationships during voluntary orienting (**Figure 1A, Active**) and passive head motion. Specifically, six mice trained in our active paradigm underwent passive head rotations where the experimenter applied an external torque to the mouse’s head, rotating it between small (∼ ±10°), intermediate (∼ ±25°), and large (∼ ±40°) angular distances **(Figure 1A, Passive VOR)**.

**#Figure 6** illustrates examples of the eye movements (magenta) evoked in these two conditions, with the corresponding head motion (blue) and gaze (yellow) motion plotted for comparison (See also **Videos S1-S4)**. As in the active orienting task, vertical eye movements were also relatively negligible during passive gaze stabilization (mean passive horizontal eye amplitude=14.0±5.4°, compared to 0.7±0.8° for passive vertical eye amplitude). In the passive condition, the application of head motion evoked an eye movement of equal and opposite amplitude, consistent with the VOR slow-phase. As a result, gaze initially remained stable **(Figure 6A, top left)**. Then, in a majority of cases (approximately 60%), this initial compensatory eye movement was followed by a subsequent rapid recentering of the eye (i.e., VOR quick-phase; **Figure 6A, bottom left)**. Importantly, the propensity of quick-phase VOR events increased with the size of the head movement, consistent with its role in recentering the eye as it approaches its mechanical limits **(Supplementary Figure 7)**. In contrast, in the active condition, the eye movement patterns showed a distinct difference, with the onset of eye movement tightly synchronized with that of head motion, regardless of whether rapid eye movement was coincident with head motion onset (**Figure 6A, top right**) or followed voluntary head motion (**Figure 6A, bottom right**). The resulting difference in head-eye coordination revealed a positive linear relationship for mean head versus eye velocity **(Figure 6B**; Active R^2^=0.05, p<0.0001), that starkly contrasted the inverse linear relationship between these two measures in the passive condition **(Figure 6B**; Passive R^2^=0.32, p<0.0001). This finding reveals a key distinction in eye dynamics between active and passive conditions. Furthermore, we found that active gaze shifts were consistently accompanied a by rapid eye redirection; in contrast, this redirection was entirely absent for approximately 40% of passive movements. We then specifically focused our analysis on the change in velocity and peak acceleration of the rapid eye movements themselves and found no difference between the active and passive conditions **(Figure 6B, Right Insets).** This is expected, given that both active saccades and reflexive VOR quick-phases are generated by the brainstem saccadic burst generator (reviewed in ^44^).

**Figure 6:**
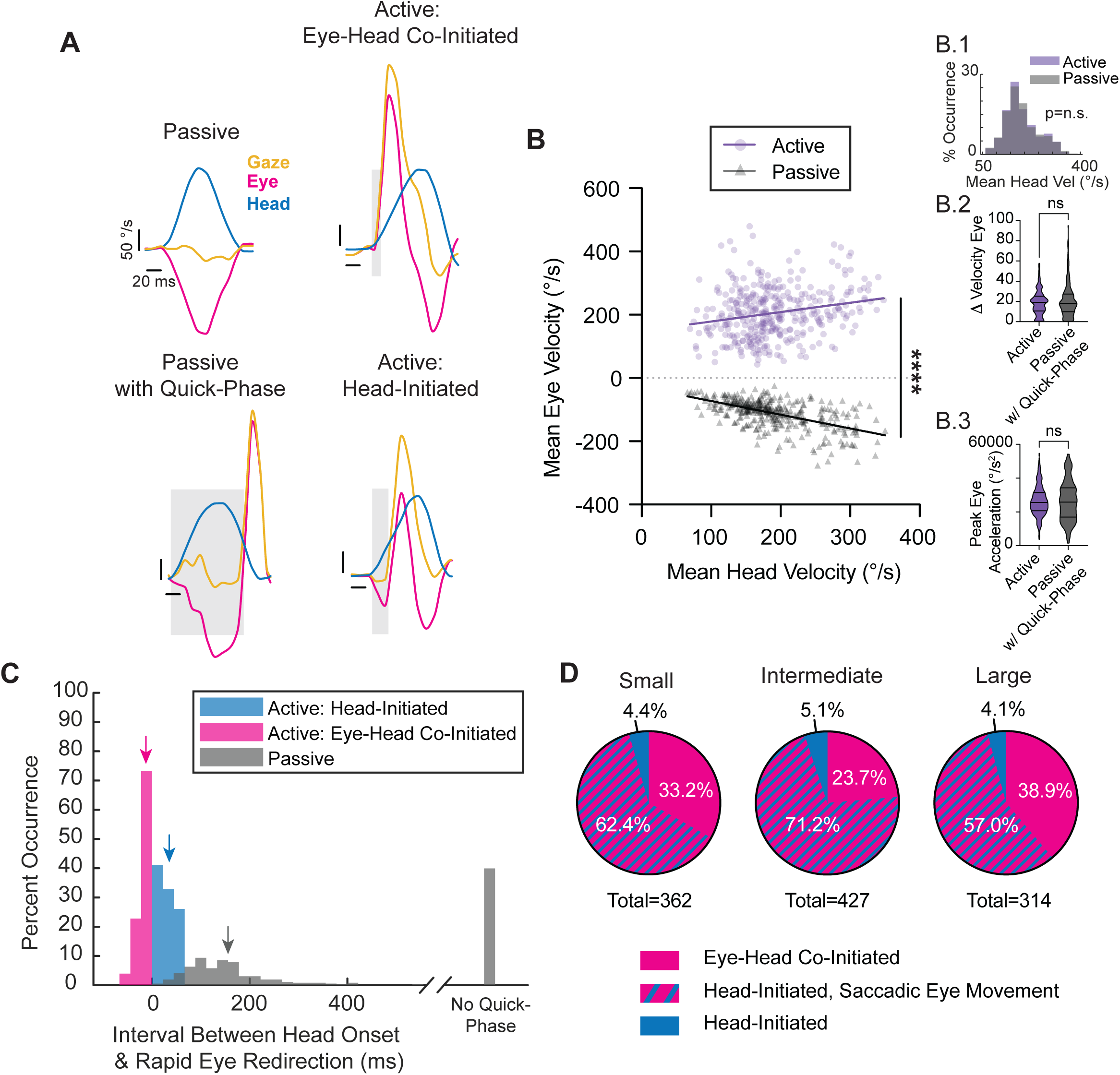
Passive VOR versus active gaze redirection. **A.** Representative velocity traces for passive gaze stabilization (top left), passive gaze stabilization with a quick-phase of nystagmus (bottom left), active, eye-head co-initiated gaze redirection (top right), and active, head-initiated gaze redirection (bottom right). Yellow = gaze. Magenta = eye. Blue = head. Vertical scale bar: 50 °/s. Horizontal scale bar: 20 milliseconds. Grey windows indicate the interval between head motion onset and rapid eye redirection. **B.** Mean eye velocities with respect to mean head velocities during head movement for active gaze shifts (purple) vs. passive gaze stabilization (grey). Active: R^2^= 0.05, slope = 0.30 (95% CI: [0.16, 0.43]), p < 0.0001, N = 361 segments. Passive: R^2^ = 0.32, slope = -0.43 (95% CI: [-0.50, -0.37]), p < 0.0001, N = 363 segments. Line comparison: F = 90.64, DFn = 1, DFd = 720, p<0.0001. Insets as follows—1: Distribution of mean head velocity between the active and passive paradigms, showing matched comparisons. Active vs. passive p = 0.8847, Wilcoxon rank-sum test; 2: Change in velocity from the start to the peak of the rapid eye movement during the active paradigm compared to the same measure during passive movements with respect to time. Welch’s t test, p = 0.1293, N = 361 active, 209 passive segmentations; 3: Peak acceleration of rapid eye movement generated during the active paradigm compared to the same measure generated during passive gaze stabilization. Welch’s t test, p = 0.9637, N = 361 active, 209 passive segmentations. **C.** Histograms of the interval between head movement onset and rapid eye redirection for head velocity–matched active: eye-head co-initiated (magenta) and active: head-initiated (blue) versus passive (grey) segments. Active eye-head co-initiated mean = -12.5 ± 14.7 milliseconds, N = 170 segments. Active head-initiated mean = 35.2 ± 17.5, N = 192 segments. Passive with quick-phase mean = 155.7 ± 83.3, N = 218 segments. Active eye-head co-initiated vs. passive with quick-phase p < 0.0001, Wilcoxon rank-sum test. Active head-initiated vs. passive with quick-phase p < 0.0001, Wilcoxon rank-sum test. Means of the respective populations indicated with an arrow. In addition, data are shown for percentage of passive movements for which no quick-phase were generated (39.9%, N = 145 segments). Histogram bin size = 22.2 milliseconds, corresponding to two video frames. **D.** Replotting of head-initiated versus eye-head co-initiated distributions shown in Figure 2 G-I, now identifying head-initiated gaze shifts demonstrating a saccadic eye movement as defined by data outside of the 6.5% overlap between active: head-initiated and passive. Small: 33.2% eye-head co-initiated, 62.4% head-initiated with saccadic eye movement, 4.4% remaining head-initiated. Intermediate: 23.7% eye-head co-initiated, 71.2% head-initiated with saccadic eye movement, 5.1% remaining head-initiated. Large: 38.9% eye-head co-initiated, 57.0% head-initiated with rapid eye redirection, 4.1% remaining head-initiated. Also see Supplementary Figure 6, Supplementary Figure 7, Videos S1-S4.

To further quantify our observations regarding the timing of rapid eye movements, we computed the interval between the onset of the head movement and the rapid eye redirection in both conditions and found a significant difference, with eye movements reaching onset in the same direction as the head markedly earlier in the active condition compared to the passive condition. This key finding underscores a fundamental difference in the dynamics of eye-head coordination across conditions. Specifically, our quantification revealed that for active gaze redirections co-initiated by eye and head movement, rapid eye motion reached threshold an average of 12.5 ± 14.7 milliseconds earlier than the head **(Figure 6C, magenta)**. This finding indicates that eye movements are not only often coincident with head movement onset but can also precede head motion onset. For active gaze redirections in which the head motion onset preceded that of eye, the latency between head and eye onset was 35.2 ± 17.5 milliseconds (**Figure 6C, blue**). In both active redirection scenarios, the eye redirected significantly faster than the 155.7 ± 83.3 millisecond delay observed before a quick-phase was initiated in the passive condition **(Figure 6C, grey)**. Indeed, only about 7% of active, head-initiated gaze redirections displayed an onset interval between head and eye movements that overlapped with those observed in the passive condition. In this context, we revisualized the gaze redirection distributions shown in Figure 2G-I to account for the rapid, “saccadic” redirection of the eye observed, faster than the eye movements in the passive conditions **(Figure 6D)**. Further, more stringent cut-off criteria would still classify the majority of rapid eye redirection as “saccadic” in nature **(Supplementary Figure 8).** Together, we identify rapid orienting eye movements in mice that serve to redirect gaze in a manner similar to cats (afoveates) and primates (foveates) **(Figure 7)**.

**Figure 7:**
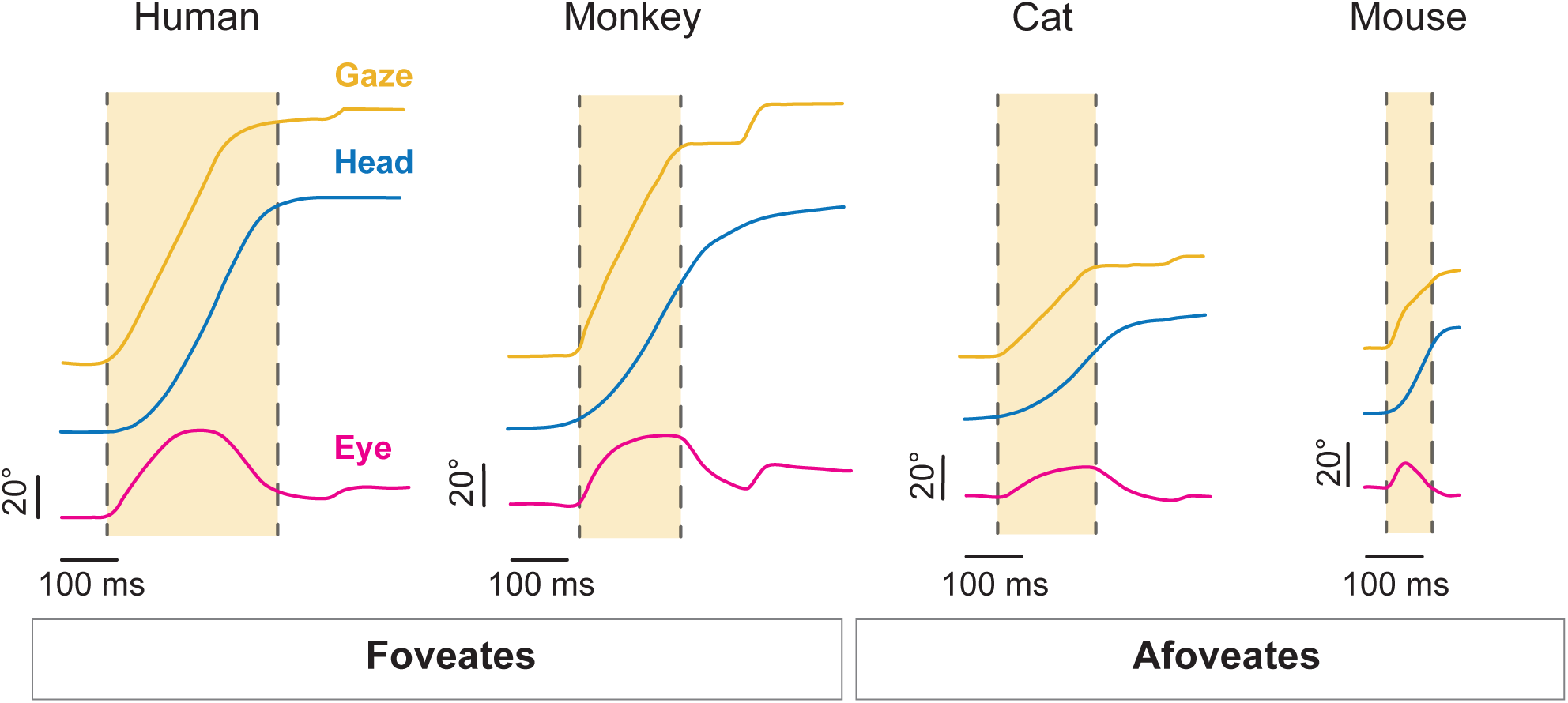
Comparison of active gaze redirection across species. Representative traces of gaze (yellow), head (blue), and eye (magenta) from human, monkey, cat, and mouse. Light yellow window indicates gaze duration. Vertical scale bar: 20°. Horizontal scale bar: 100 milliseconds. Human, monkey, and cat traces adapted with permission from Guitton 1992 ^38^.

## DISCUSSION

Our central finding is that the eye and head movements of mice are coordinated in a manner similar to foveate species during active orienting. Specifically, we demonstrate that mice, despite being afoveate, generate active saccadic eye movements that are tightly synchronized with active orienting head movements. Importantly, these active saccadic eye movements occur at markedly shorter latencies than the reflexive quick-phase eye movements evoked by comparable passive head rotations. To our knowledge, this is the first study to both directly measure eye versus head motion onset during voluntary orienting in mice and compare the relative latencies with those evoked by passive head motion. These active saccadic movements were directed in the same direction as the active head movement, and their onset was tightly coupled with head motion, with a substantial proportion leading head motion onset. Overall, the eye and head movements produced during active orienting followed relationships comparable to those observed in other species, including monkeys and humans. Thus, taken together, our present results suggest that a common neural strategy for eye-head coordination is conserved across vertebrate species.

### Saccades generated by mice

As locomotion across species evolved, changes in anatomy and locomotion strategy posed challenges for maintaining stable vision. This is because our eyes are attached to our heads; with every head movement, retinal slip would occur, preventing us from maintaining stable images of the visual world. As such, species developed the gaze-stabilizing VOR consisting of two parts: i) a slow-phase that counteracts head movement to stabilize gaze, and ii) a fast phase that rapidly moves the eye at high speeds in the direction of head velocity, preventing any attempt to move the eyes beyond the oculomotor range^44, 45^. As is the case for nearly all vertebrate species^16^, mice generate robust VOR eye movements that stabilize gaze during both passively applied head motion (e.g., horizontal head rotations^46–48^ and head tilts^49–51^) as well as during head motion in freely moving rodents^16–20^. With the evolution of the high-acuity fovea, the need for an independent eye movement system to focus on specific objects arose in foveate species, leading to the development of saccadic eye movements^44, 45^. Accordingly, in foveates, eye movements can be divided into two categories: those that stabilize gaze via passive reflexes, and those that are voluntary and actively target gaze to a new point of interest.

During natural behavior, afoveate animals such as mice use a “saccade and fixate” system to visually explore their surroundings^6, 11^. However, although the quick eye movements seen in mice are often referred to as “saccades” in the literature, one prevailing view has been that they are not actively generated to target a new object of interest, as seen in foveates. Instead, these rapid eye movements were generally thought to be reflexively driven by active orienting head movements that redirect their gaze (e.g.,^6, 11, 15, 16^). In this view, when mice shift their gaze, they make an active head movement in the direction of the object. In turn, the resulting vestibular stimulation reflexively evokes an oppositely-directed slow-phase VOR eye movement and, in some cases, a subsequent VOR quick-phase that rapidly redirects the eye in same direction as head motion. Recent studies in mice have demonstrated that, even after accounting for head rotations, horizontal eye position can exhibit a strong correlation between both eyes^18^. However, it had remained unclear whether these correlations result from eye movements resetting independently of head movements, as suggested in ^52^, or if they are the result of active gaze shifts. Here we establish that (i) the onset of these active saccadic movements is closely coupled with head motion, with a substantial proportion leading the onset of head motion, and (ii) these active saccadic eye movements occur at significantly shorter latencies than the reflexive quick-phase eye movements evoked by comparable passive head rotations, demonstrating context-dependent differences in eye movement generation that have also been reported in other vertebrates^53, 54^. Thus, our present findings provide direct evidence that the rapid eye movements made by mice during gaze redirection are actively generated.

Our finding that rapid eye movements during active orienting occur at shorter latencies than reflexive quick-phases driven by vestibular feedback is interesting to consider in light of prior head-restrained studies on mouse eye movements. In particular, head-restrained mice can generate rapid eye movements resembling saccades, although whether those movements were produced voluntarily remained a question^18, 23, 37, 55–58^. Importantly, the presence of a head restraint in these prior studies inherently eliminated vestibular feedback, demonstrating that mice can make rapid eye movements in the absence of vestibular input. Here by studying gaze control under more naturalistic conditions, our work goes beyond these prior findings to reveal that rapid eye movements during active orienting are tightly coupled to head motion during voluntary orienting behaviors, highlighting the role of the eye in actively coordinating gaze. Further, our results demonstrate that mice readily engage in goal-directed orienting between waterspouts, consistent with the broader trend that mice can rapidly learn directional, water-rewarded tasks that parallel primate ‘center-out’ frameworks^59^. Given this task structure, we hypothesize that the relative proportion of head-initiated versus eye-head co-initiated orienting movements is not fixed but instead depends on behavioral context. Specifically, this distribution may shift towards a higher prevalence of eye-head co-initiated during more exploratory behaviors or in response to unexpected stimuli. This prediction is consistent with primate studies showing that head-free gaze shifts to sudden or unpredictable targets are typically initiated by a saccade, with task demands strongly shaping eye-head coordination strategies^4, 60–62^.

### The coordination of eye and head movements during active orienting

Many animals use a combination of eye and head movements to rapidly orient their visual focus (gaze), which are commonly referred to as gaze shifts. Previous studies in primates and cats have shown that a shared eye-head gaze controller drives both eye and head movements to achieve accurate gaze redirection^45, 63^. This mechanism generates a dynamic gaze feedback signal to drive both the head and eye premotor circuits until gaze lands on the intended target and gaze error is nulled^60, 64–66^. Thus, instead of enforcing a specific eye or head trajectory, it minimizes gaze errors^30, 67, 68^, similar to the concept of optimal feedback control (for review, see ^69–71^). While the underlying principles of gaze control apply to humans, rhesus monkeys, and cats, there are differences in how each species uses eye and head motion.

Specifically, these differences are linked to the range of motion of a given species’ eyes, known as the oculomotor range (reviewed in ^38^). For example, humans and monkeys have a relatively large oculomotor range of about ± 55 degrees, such that they rely more on eye movement to shift gaze^38, 72^. In contrast, cats have a more limited oculomotor range of about ± 25 degrees, necessitating more head motion for gaze redirection, especially for larger shifts^38^. As a result, primates display less stereotyped head motion than cats (reviewed in ^64^) but still must generate head motion for larger gaze shifts^7^. Here we present evidence that mice, like cats, monkeys, and humans, also generate active eye movements when orienting. Although our orienting behavioral paradigm rewarded solely based on head movement, we found that the eyes actively contribute to gaze redirection, reinforcing a natural eye-head coupling in mice during everyday orientation. Notably, mice have a relatively small oculomotor range of about ± 20 degrees^18, 56, 72^. Yet we found that the coordinated eye-head gaze shifts generated by mice follow the same patterns (i.e., main-sequence relationships, dependency of head motion on initial eye position in orbit) as those observed in primates and cats^4, 7, 32, 73^.

In cats and primates, the SC is known to control active orienting by generating integrated motor commands that couple saccadic eye movements with head movements, thereby coordinating gaze shifts during voluntary behavior^67^. Consistent with this role, stimulation of the SC in these species evokes coordinated eye-head gaze shifts (cats^28, 74, 75^ and monkeys^8^). We propose that the SC may serve a similar role in mice. While early SC stimulation studies in head-unrestrained rodents reported orienting movements (mice^31, 76, 77^, rats^78^), these studies did not focus on eye movements. More recent work in head-restrained mice demonstrated a tight temporal association between eye movement onset and head torque against the restraint—interpreted as an attempted head movement—following SC stimulation^23^. A follow-up study in head-unrestrained mice recorded both evoked eye and head motion and found a similarly tight coupling^43^. Although these studies did not explicitly report the latency between evoked eye and head motion, both report values suggesting delays under 30 milliseconds^23, 43^.

Our present findings extend this literature by providing the first direct evidence that during voluntary, goal-directed orienting, freely moving mice generate tightly coordinated eye and head movements with similarly short latencies. Importantly, these movements follow an active gaze redirection strategy in which eye motion onset often reached threshold prior to head motion—indicating that eye movements are not merely reflexive but can actively lead the orienting response. This pattern is consistent with previous reports that saccades in head-restrained mice are generally accompanied by attempted head movements ^18, 23^, and that when saccades are evoked by sensory stimuli, eye motion can begin before head motion ^23^. Thus, taken together, these observations support the idea that mice employ conserved visuomotor strategies shared with primates and other mammals, where the eye’s lower inertia can enable faster initiation to reorient gaze while the head with its greater inertia follows. Such strategies likely optimize visual processing in dynamic environments ^38^. Although our study does not directly test the role of the SC, the behavioral pattern we report is consistent with prior findings from SC stimulation studies in mice. Accordingly, we speculate that the SC likely contributes to coordinating eye-head coupling during natural orienting in mice, as it does in other species. Future work should aim to identify the specific circuits and computations through which the mouse SC—and potentially other brainstem and cortical structures—coordinate goal-directed and reflexive eye-head movements in complex environments.

### Functional implications of active saccadic eye movement in mice

Our finding that mice coordinate their eye and head movements during active orienting in a manner similar to that used by foveate species raises the question of why it would be functionally advantageous for mice to actively move their eyes. One obvious potential advantage is that it allows mice to target objects of interest more precisely. While humans and monkeys possess a foveal specialization in their retina (a small central area with high photoreceptor density for optimal visual acuity), cats also have a retinal specialization, although not a traditional fovea. Instead, cats have a slit-shaped visual streak with a relatively higher photoreceptor density compared to the surrounding retina^79–81^. Consequently, it makes sense that cats, like humans and monkeys, would generate voluntary eye movements to accurately place their eyes on a target of interest. Similarly, while mice lack fovea, their retinal anatomy is also not entirely uniform. The central region of the mouse retina has a higher photoreceptor cell density than the periphery, albeit not as pronounced as in primates^82, 83^. This non-uniform distribution is further evident within the retinal ganglion cell (RGC) layer^84^, thereby enabling mice to sample their frontal visual fields more effectively; this correlates with a fovea-like representation within their visual cortex^85^. Further, it has been shown that neurons within mouse V1 preferentially respond to gaze redirecting rather than compensatory movements^86^. We speculate that, even though mice do not have a conventional fovea, the variation in photoreceptor distribution and central retinal anatomy allow for strategic, preferential focus on objects within the frontal visual field, as this would be evolutionarily advantageous in both prey and predator scenarios.

Another, not mutually exclusive, advantage of mice generating active eye movements is that this ability could allow them to achieve binocular viewing of a target of interest. Unlike primates with front-facing eyes, mice have lateral-facing eyes that provide a broad field of view for detecting potential predators (approximately 280° ^6, 15, 16, 23, 24, 87, 88^). This wide field of view comes at the expense of a smaller overlapping binocular field (about 40° versus 135° in humans^89^). Nevertheless, neurons in the mouse visual cortex do encode binocular disparities^90, 91^, suggesting that optimizing binocular alignment may have functional benefits (see also ^18^). Furthermore, when mice are locating prey within a visual scene, the prey image consistently falls within a stabilized portion of the visual field that coincides with the region of minimal optic flow within the visual field, thereby resulting in minimal motion-induced image blur^92^. Similar stabilization of visual input, supported by coordinated eye-head movements, has also been reported in other species, including ferrets^93^. Interestingly, the population of RGCs that makes up the retinal population within this region of minimal optic flow (Alpha-ON) are non-uniformly distributed and permit enhanced visual sampling within certain retinal regions^84^, as described above. Such stabilization would in fact need to be properly integrated with an ability to shift the axis of visual gaze to achieve the intended behavior during active prey location. Stereopsis, the ability to perceive depth through binocular vision, was originally thought to be exclusive to mammals with front-facing eyes, primarily for prey detection^94^. However, it has now been observed in various vertebrates and invertebrates, including those with laterally-placed eyes like mice, and has been proposed to play a more general role in essential behaviors such as navigation, predator avoidance, and prey capture^16, 92, 95^. Taken together, our present findings support the view that eye movements have a conserved role in active orienting across species, including mice.

## Supporting information

Video S1: Active Head Initiated

Video S2: Active Eye-Head Co-Initiated

Video S3: Passive VOR

Video S4: Passive with Quick Phase

## ACKNOWLEDGEMENTS

The authors are grateful to O. E. L. Brown, M. Green, X. Li, Y. Liu, T. Niebur, O. Stanley, S. Somanathan, Y.S. Sun, K.P. Wiboonsaksakul, and O. Zobeiri for their experimental contributions, as well as all members of the Cullen Lab for their thoughtful feedback throughout the creation of this manuscript. This work was supported by the National Institutes of Health grants K12-GM123914 (BMV) and 1U01-NS111695 (KEC et al.).

## AUTHOR CONTRIBUTIONS

BMV, HHVC, and KEC designed the experimental study. BMV and HHVC performed surgical procedures. BMV oversaw animal training and husbandry. DCR created and optimized the video oculography interface. BMV performed all behavioral acquisitions and analyses. BMV and KEC wrote the manuscript with input and approval from all authors. This work was supervised by KEC.

## MATERIALS & METHODS

### Experimental Model and Subject Details

Nine adult (8 months old) male C57BL/6J mice (The Jackson Laboratory) were used in this study. All animals used in this study were bred and maintained on site. Mice were housed in standard cages in a temperature and humidity-controlled animal facility on a traditional 4.5% fat diet *ad libitum*. Animals had *ad libitum* access to water (approximately 500 mL) when not engaged in the active training paradigm (see Method Details: Active and Passive Behavioral Paradigms). Animals used in this study were individually housed post-surgery (see Method Details: Headpost and Camera Mount Implantation). Animals were on a 12:12 h light: dark cycle and tested during the light phase. All animal husbandry and procedures were carried out with ethical approval and oversight from the Animal Care and Use Committee at Johns Hopkins University, National Institutes of Health Guide for the Care and Use of Laboratory Animals, and in accordance with ARRIVE Guidelines^96^. We have complied with all relevant ethical regulations for animal use.

### Headpost and Camera Mount Implantation

Mice were anesthetized via isoflurane inhalation (1.5-2.5% isoflurane in O_2_) and secured in a stereotaxic surgical frame. A midline incision was performed to secure the skull, followed by the addition of topical bupivacaine for pain management. A thin layer of dental adhesive (Kerr, 314614) was applied to the skull, followed by the placement of a custom-built headpost over bregma. The camera mount was a 4x4 receptacle connector (approximately 3x3 mm) placed immediately anterior to the headpost on the surface of the skull and secured with dental acrylic. Skin was re-adhered around the implant with Vetbond (3M, 014006) and topical analgesics and antibiotics was administered at the incision borders. Animals were monitored for a minimum of 24 hours post-op before any experimentation.

### Active and Passive Behavioral Paradigms

For the active paradigm, mice were restricted from receiving water for 48 hours prior to the first day of training. Mice remained water-restricted throughout the study with the exception of waterspout reward delivery and 100 mg of H_2_O gel following a training session (details in the forthcoming paragraphs).

Animals were placed in a custom-printed tube at the center of a turntable. Headposts attached to the mice were connected to custom-printed arms attached to a low friction potentiometer that measured the change in horizontal head position as a measure of voltage change. Mice were allowed to settle into a comfortable resting head pitch prior to fixation to the potentiometer. Movement was largely restricted to the horizontal (yaw) axis, comparable to the devices used in prior studies of primate gaze shifts (e.g., ^4, 7, 35^).

Two waterspouts were placed at approximately ± 10°, ± 25°, or ± 40° from center to evoke different sized orienting movements. The order of spout distance was randomized between mice and session. Head position was monitored in real time using a QNX-based data acquisition system (REX) and SpikeGLX (Janelia) **(Schematic: Supplementary Figure 1A)**. Animals were trained to move their head from spout A directly to spout B at the sound of a randomized tone (50 ms, 80 dB, 4.1 kHz, interval range=2.5-7 seconds) (and vice versa, spout B to A) at increasingly spaced distances to receive a water reward. No reward was given for incomplete movements; for example, pausing between the spouts or failure to move within the reward window set around tone onset. Spout position corresponded to a 10° reward window, in which a reward of approximately 40 µL H_2_O was delivered once the mouse’s head entered the positional window **(Supplementary Figure 1A).** For the purposes of this study, success/failure was defined by heading towards the reward (ie. the head entering the reward window) and not necessarily consumption of the reward, though H_2_O consumption was confirmed by the experimenter at multiple points throughout the training session by assessing the presence or absence of water accumulation.

Training began 2 weeks prior to acquisition, consisting of daily 1-hour training sessions to acclimate the mouse to tone and water delivery. Study acquisitions consisted of three 10-minute trials with a 3-minute inter-trial interval. The timing criterion for successful completion of the task was purposely relaxed to minimize the risk of repetitive movement patterns, thus encouraging more naturalistic behavior while still ensuring that mice were engaged in the active orienting task. Mice performed an average of 120 attempted head movements in a 10-minute session. Successful orienting movements were included if (1) the mouse’s head entered the reward window, (2) eye movements were successfully tracked for the duration of a head movement without interruption (e.g., blinking), and (3) the resulting eye, head, and summated gaze signals all reached minimum onset threshold in the intended direction (see Methods: Eye, Head, and Gaze Analysis). Well-trained mice indeed anticipated the tone, but no difference between strategy (head-initiated vs. eye-head co-initiated) was observed **(Supplementary Figure 1C)**. As such, all completed head movements, regardless of tone anticipation, were quantified in this study. For the passive VOR paradigm, no water reward was administered, and instead the experimenter used their hand to apply an external torque to the mouse’s head. The head was moved back and forth from positions approximating ± 10°, ± 25°, and ± 40° from center with position measured by the potentiometer. For both the active and passive behavioral paradigms, an opaque surround separated the mouse from the experimenter.

Eye and head movements were also recorded during two additional active paradigms. First, we restricted the mouse’s body to prevent head movement about the lumbar column during our active behavioral paradigm **(Supplementary Figure 2)**. This paradigm allowed us to focus exclusively on head movements generated by the head-neck ensemble, isolating them from any contributions by body motion. Specifically, mice were body-wrapped about the thoracic and lumbar regions of the spine and placed back in the custom-printed tube at the center of the turntable (termed additional lumbar restriction). They then repeated the active paradigm while body-restricted, as described above. Second, to confirm that our active setup did not alter the timing of eye versus head movements, mice made orienting movements in an additional paradigm which allowed completely free head motion (‘Head-Free’). We tracked their head movements using a miniature inertial measurement unit (IMU) attached to the headpost **(Supplementary Figure 3)**, and a body restriction collar was put in place to prevent the mouse from exiting the tube during the head-free assessment. In this second active experiment, mice again performed the active paradigm as described above (‘Yaw-Restricted’) with waterspouts placed at ± 25° while the IMU captured the angular rate and linear acceleration of the head. Mice then performed the active paradigm in a head-free manner, orienting between the two spouts while not attached to the potentiometer.

### Eye and Head Tracking

To track the position of the pupil, video oculography software (VOG) was used in conjunction with a small camera (Arducam, B0066-02) with a frame rate of 90 FPS placed in front of the mouse’s eye. The camera was held in place by a custom 3D printed camera holder connected to the mouse headpost. This allowed the camera to move with the head while maintaining fixation on the mouse’s eye. VOG was captured onto the Raspberry Pi 4 (RAS-4-4G). The OpenCV (https://opencv.org/) image processing library was used to threshold the eye image and detect the dark pupil via the findContours function. The centroid of the pupil contour was computed using the moments function. This permitted resolution beyond one pixel. The frame-to-frame pupil tracking variability was approximately ± 0.19°. To time-align the acquired pupil position dataset with the head position data, a frame sync pulse was delivered from the Raspberry Pi 4 to SpikeGLX for *post hoc* synchronization by aligning each VOG data sample with its corresponding sync pulse edge. Oscilloscope visualization confirmed sync signal delivery with a variability of 0-40 µs, below the millisecond estimations used in subsequent analyses. Head yaw angle was measured as a voltage from the potentiometer, sampled at 5 kHz into SpikeGLX and converted to degrees *post hoc*. For experiments using an IMU, data was time-aligned using a data sync pulse delivered from the IMU to SpikeGLX. Head position was then derived from obtained head velocity data and compared in the yaw-restricted versus head-free conditions. Both horizontal head and eye position were imported into Matlab (Mathworks) and head position data was down-sampled from 5 kHz to 90 Hz to match the native sampling of eye position for subsequent analysis (See **Supplementary Figure 1D, E** for representative VOG processing steps). This permitted positional detection limited to the resolution of the camera’s sampling period.

### Eye, Head and Gaze Analysis

During VOR in light, opposing head and eye movements at 1-2 Hz yield a gain near unity (e.g., ^97, 98^). To calibrate eye positions to degrees, mice underwent ∼1 Hz (range: 1.0-1.6 Hz) sinusoidal yaw rotations under light conditions while head-fixed, with angular head velocity recorded using a gyroscope. Movements were timed to a metronome cue to minimize inter-trial variability. Calibration peak velocities ranged from 54-75°/s (mean: 62°/s). The average number of movements and subsequent cycles was approximately 34 and 18, respectively (movement range=22-64 movements; cycle range=11-32 cycles). Gyroscope data were captured in SpikeGLX, and pupil positions were recorded with Raspberry Pi VOG, synchronized to SpikeGLX (See Method Details: Eye and Head Tracking). Eye velocity, derived from synchronized pupil positional data, was scaled to match head-eye gain to 1, representing degrees-per-second.

Down-sampled head and native eye position data acquired from both the active and passive behavioral paradigms were summed to calculate gaze. Head, eye, and gaze position data was then digitally low-pass-filtered (Butterworth, cut-off frequency = 40 Hz). Identical filtering of both the active and passive behavioral data allowed direct comparisons between eye and head movement latencies both within and across conditions (see ^99, 100^). All three position signals were differentiated to obtain their respective velocities. For each successful trial, the onsets and offsets of head and gaze redirection were determined using a 25°/s threshold criterion used in prior studies of primate gaze shifts (e.g.,^4, 7^). Likewise, the onsets and offsets of rapid eye redirection made in the direction of head motion were determined using a 25°/s threshold criterion, relative to any compensatory slow-phase eye movement. During active gaze redirection, gaze shifts were classified as “Head-Initiated” when head velocity reached its onset threshold before both eye and gaze movement. They are classified as “Eye-Head Co-Initiated” when eye and gaze velocity reached onset threshold at or before the onset of head motion. Metrics of head, gaze, and eye position and velocity were then computed within each of these onset and offset windows. For passive head movements, the onsets and offsets of head, gaze, and eye movement were computed using the same approach. All analyses were performed on the down-sampled head and native eye data (90 Hz). Data was resampled to 1 kHz after analysis for visualization.

### Statistics and Reproducibility

Statistical analyses were performed using Matlab. Histogram comparisons were performed using a two-sided Wilcoxon rank sum test. Means shown as mean ± standard deviation unless otherwise indicated. Linear and nonlinear fits were performed in Prism 9 (GraphPad). Simple linear regressions used non-zero significance without forcing through origin. Exponential fits were computed using GraphPad’s “one phase association” equation: Y = Y0 + (Plateau - Y0) * (1 - exp(-K * x). For analyses performed on individual movements, the unit of analysis was a single active orienting movement or applied head movement; for each condition, figure legends report both the number of movements (i.e. “segments”) analyzed and the number of mice from which those movements were obtained (e.g., “N = X segments across Y mice”). All reported effects were observed across multiple mice and were reproducible across animals.

## DATA AVAILABILITY

Requests for further information should be directed to the lead contact, Kathleen Cullen. Data supporting the findings of this study are available in FigShare with the identifier https://doi.org/10.6084/m9.figshare.31686991. Any additional information required to reanalyze the data reported in this paper is available from the lead contact upon reasonable request.

## CODE AVAILABILITY

Custom MATLAB scripts were used to generate the figures and is available from the corresponding author upon reasonable request.

