## Supplementary Data for "Eye-head coordination during goal-directed orienting in mice"

**Supplementary Information**

Supplementary Figures 1-8
Video S1. Eye and head position during Active: Head-initiated gaze redirection, Related to Figure 2, Figure 6, Figure S1
Video S2. Eye and head position during Active: Eye-Head Co-Initiated gaze redirection, Related to Figure 2, Figure 6, Figure S1
Video S3. Eye and head position during Passive VOR, Related to Figure 6
Video S4. Eye and head position during Passive with Quick-Phase, Related to Figure 6

### Supplementary Figure 1

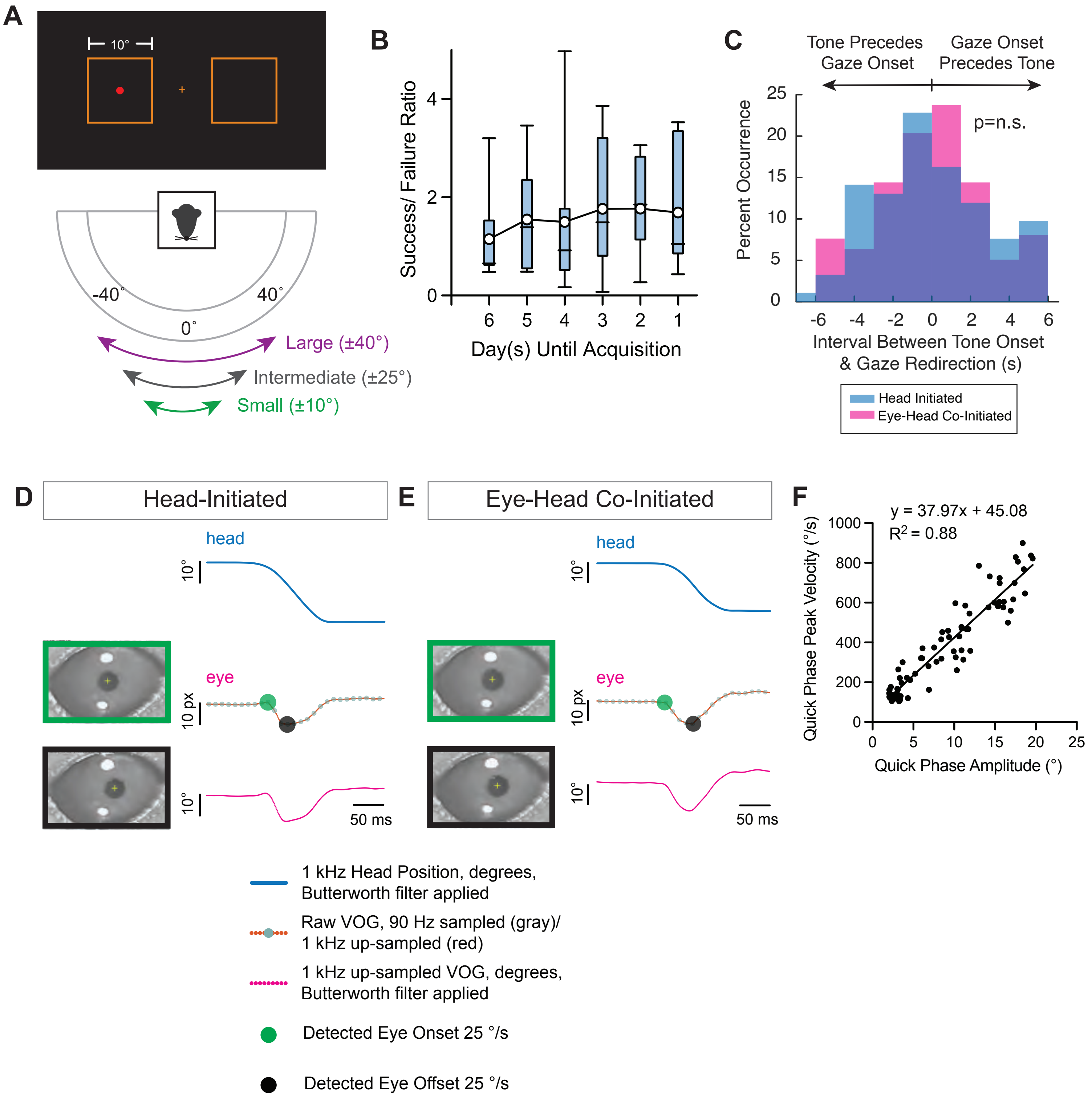

**Supplementary Figure 1: Additional behavioral and training metrics, Related to Figure 1. A.** Schematic of the potentiometer output within the QNX-based data acquisition system (REX). Red dot indicates the position of the mouse's head within a reward window corresponding to a 10° range about spout position. Cross corresponds to the center between two spouts. Figure 1B schematic shown for spout placement reference. **B.** Box-and-whisker plot of success/failure ratio with respect to the days leading up to a formal behavioral acquisition for a subset of experimental mice. Mean trendline overlaid. N=7 mice; Means ± SEM: Day 6 = 1.14 ± 0.37; Day 5 = 1.55 ± 0.43; Day 4 = 1.50 ± 0.62; Day 3 = 1.77 ± 0.52; Day 2 = 1.77 ± 0.38; Day 1 = 1.69 ± 0.48. **C.** Distribution of the interval between tone onset and gaze redirection for successfully attempted gaze redirections. Head-initiated (blue): Mean = -0.0041 ± 3.004 seconds, N = 236 gaze segments from 3 mice. Eye-Head Co-Initiated (magenta): Mean = -0.0007 ± 2.724 seconds, N = 92 gaze segments from 3 mice. Head-initiated vs. eye-head co-initiated p = 0.8453, Wilcoxon rank-sum test. Top arrows indicate portion of the distribution in which tone delivery preceded gaze onset (left) or when gaze onset preceded tone delivery (i.e. anticipation) (right). No difference in timing re. gaze onset in comparison with Figures 2G-I was observed when stratified by anticipation status. **D, E.** Head position (top, blue) resampled to 1 kHz with the Butterworth filter applied, along with the processing steps of the eye position data for a (D) head-initiated and (E) eye-head co-initiated gaze redirection; Middle: raw VOG samples at 90 Hz, demonstrating an ~11 millisecond spacing between data acquisition points (gray dots), along with the VOG data up-sampled to 1 kHz (red dots); Bottom: up-sampled VOG data in degrees with the Butterworth filter applied (magenta). Green and black dots identify the onset (green) and offset (black) of the eye redirection as calculated by a 25 °/s threshold criterion derived from position. Still frames of the eye immediately preceding the detected onsets and offsets shown. Also see Video S1 and Video S2 for video segments corresponding to Supplementary Figure 1 D & E. **F.** Quantification of the main-sequence relationship of quick-phase eye movements during VOR calibration. Linear Regression, $R^2$ = 0.88, $p < 0.0001$, best-fit y = 37.97 ± 45.08; n=77 quick-phases, range = 6-13 quick-phases across 9 mice.

**Supplementary Figure 2**

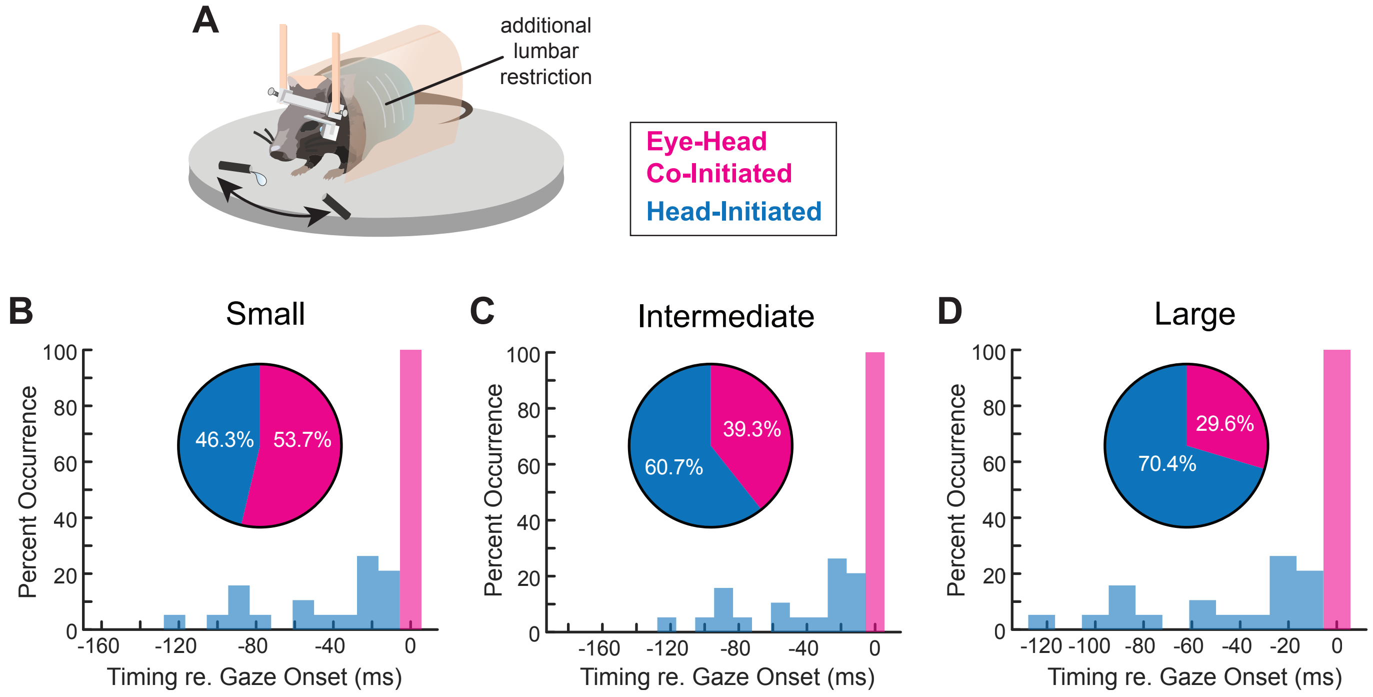


**Supplementary Figure 2: Distribution of gaze redirections with additional body restriction, Related to Figure 2. A.** Mice received additional body restriction prior to being placed inside the tube for performance of the active behavioral paradigm. **B-D.** Distribution (insets) and timing with respect to gaze onset of small, intermediate, and large head-initiated and eye-head co-initiated orienting movements during task performance with additional body restriction. Blue= head-initiated. Magenta= eye-head co-initiated. **B.** Small inset: 46.3% head-initiated versus 53.7% eye-head co-initiated. Mean latency head: -32.7 ± 29.1 milliseconds. Mean latency eye: -0.79 ± 3.7 milliseconds. Head-initiated v. eye-head co-initiated $p < 0.0001$. N = 162 segments across 3 mice. **C.** Intermediate inset: 60.7% head-initiated versus 39.3% eye-head co-initiated. Mean latency head: -43.5 ± 37.0 milliseconds. Mean latency eye: -0.01 ± 0.06 milliseconds. Head-initiated v. eye-head co-initiated $p < 0.0001$, N = 89 segments across 3 mice. **D.** Large inset: 70.4% head-initiated versus 29.6% eye-head co-initiated. Mean latency head: -47.9 ± 45.9 milliseconds. Mean latency eye: 0.0 ± 0.0 milliseconds. Head-initiated v. eye-head co-initiated $p < 0.0001$, N = 27 segments across 3 mice. Histogram statistical comparisons performed using Wilcoxon rank-sum test. Latencies shown as mean ± SD. Histogram bin size = 11.1 milliseconds, corresponding to a 90 Hz sampling rate, or 1 video frame. Comparison of the distribution of head-initiated vs. eye-head co-initiated with additional body restriction compared to Main Figure 2 data as follows: Small: $\chi^2 = 18.93$, dof = 1, $p < 0.0001$; Intermediate: $\chi^2 = 8.53$, dof = 1, $p = 0.0035$; Large: $\chi^2 = 0.55$, dof = 1, $p = 0.459$. Chi-square independence test.

**Supplementary Figure 3**

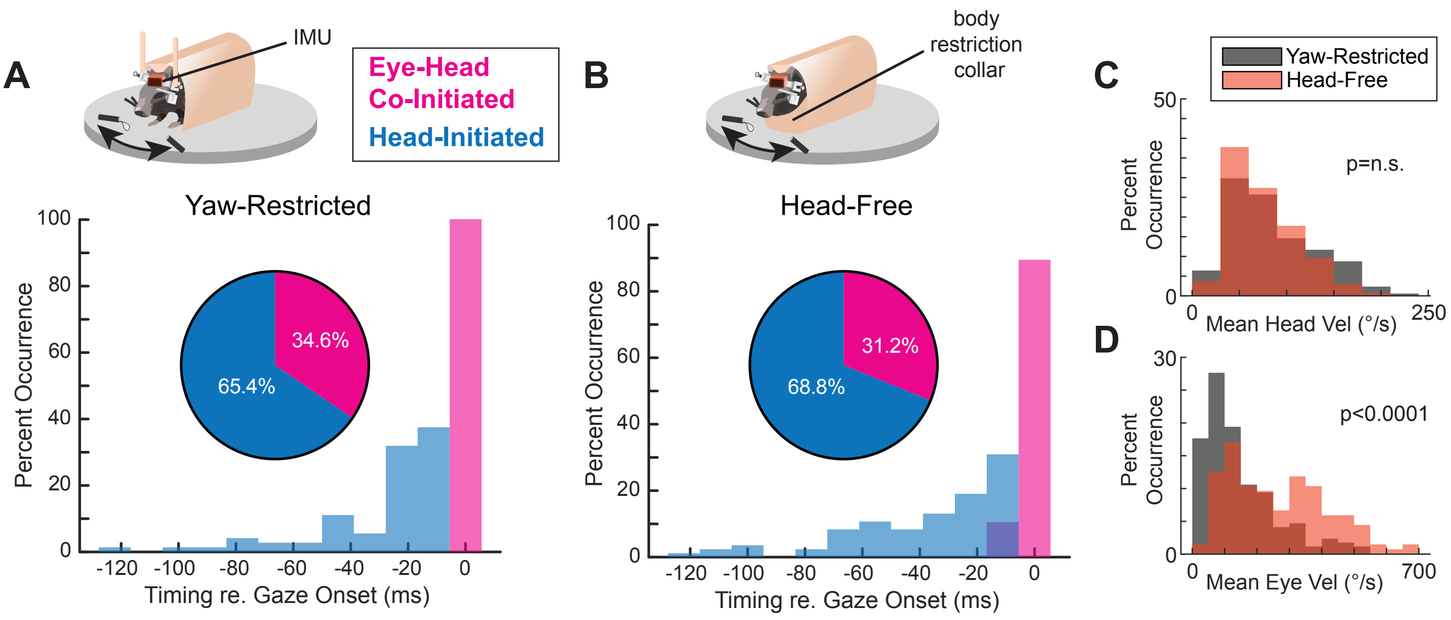


**Supplementary Figure 3: Distribution of gaze redirections during yaw-restricted and head-free active behavior, Related to Figure 2. A, B.** Distribution (insets) and timing with respect to gaze onset of yaw-restricted and head-free orienting movements. Blue = head-initiated. Magenta = eye-head co-initiated. **A.** Yaw-restricted distribution: 65.4% head-initiated versus 34.6% eye-head co-initiated. Mean latency head: -29.0 ± 23.8 milliseconds. Mean latency eye: 0.0 ± 0.0 milliseconds. Head-initiated v. eye-head co-initiated $p < 0.0001$, N = 110 segments across 3 mice. **B.** Head-free distribution: 68.8% head-initiated versus 31.2% eye-head co-initiated. Mean latency head: -36.7 ± 27.8 milliseconds. Mean latency eye: -1.2 ± 3.4 milliseconds. Head-initiated v. eye-head co-initiated $p < 0.0001$, N = 122 segments across 3 mice. Histogram statistical comparisons performed using Wilcoxon rank-sum test. Histogram bin size = 11.1 milliseconds, corresponding to a 90 Hz sampling rate, or 1 video frame. Latencies shown as mean ± SD. Comparison of the distribution of head-initiated vs. eye-head co-initiated during yaw-restricted compared to head free: $\chi^2 = 0.169$, dof = 1, $p = 0.681$, suggesting that the distributions are independent of experimental condition. Chi-square independence test. Schematics of the behavioral acquisition setup for yaw-restricted (A) and head-free (B) conditions shown. In both conditions, an IMU was attached to the mouse's headpost to measure the velocity and acceleration of the head. Spouts were positioned ± 25° for both yaw-restricted and head-free. In (A), mice perform the active behavioral paradigm as described, which consists of mice being attached to a potentiometer. In (B), a body restriction collar is put in place, the potentiometer is released, and mice repeat the active paradigm. **C.** Distribution of mean head velocity between the yaw-restricted and head-free paradigms. $p = 0.27$, Wilcoxon rank-sum test. **D.** Distribution of mean eye velocity between the yaw-restricted and head-free paradigms. $p < 0.0001$, Wilcoxon rank-sum test.

**Supplementary Figure 4**

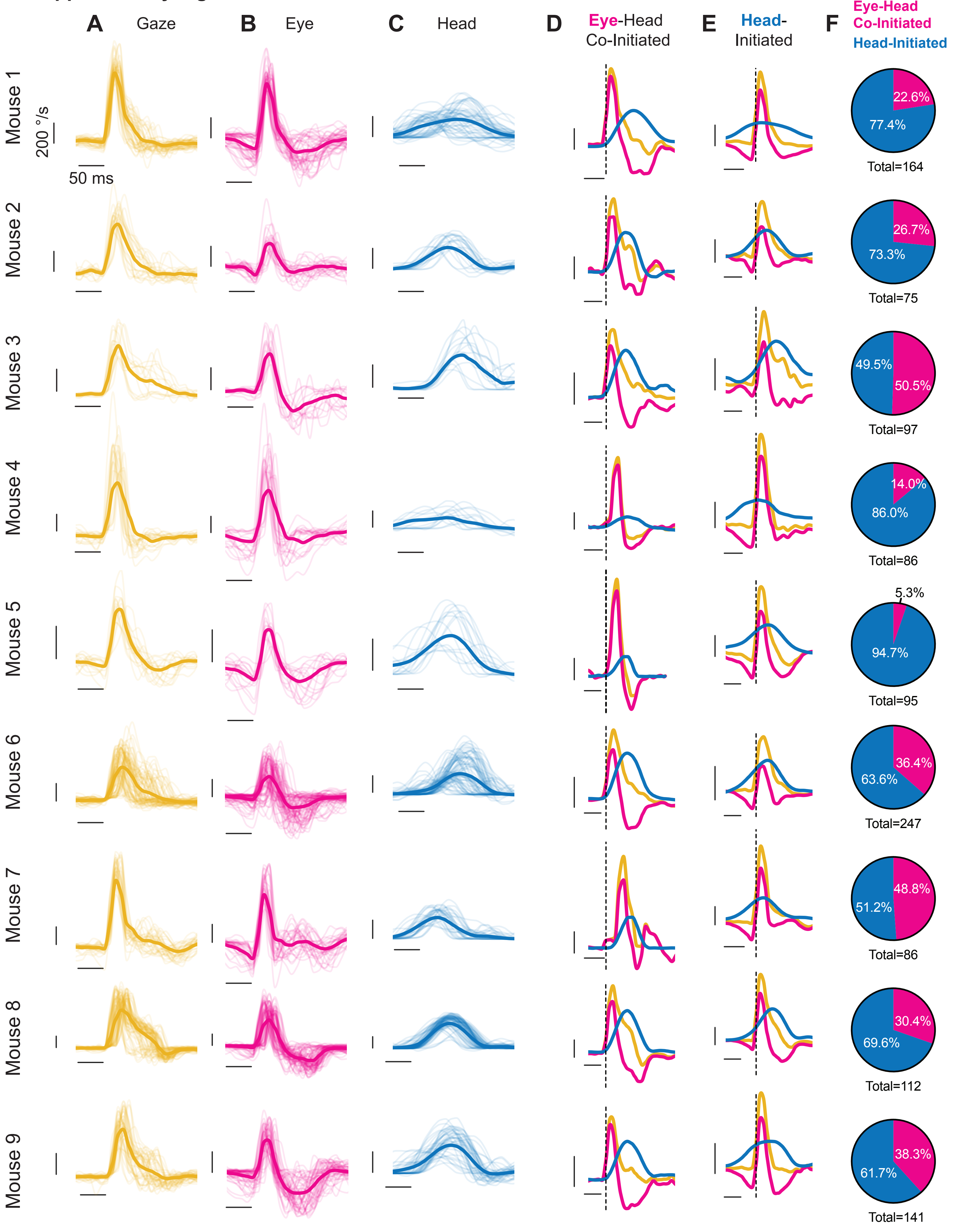

**Supplementary Figure 4: Head, eye, and gaze redirection velocities per mouse, Related to Figure 3. A-C.** Gaze (A), eye (B), and head (C) redirections as shown as velocity traces of large orienting movements for all 9 mice in a given direction. Eye and head onsets aligned to gaze onset. Yellow = gaze. Magenta = eye. Blue = head. Individual traces shown as thin lines. Means of respective signal as thick line. **D, E.** Means of gaze, eye, and head velocities from A. now shown as eye-head co-initiated (B), and head-initiated (C) orienting movements per mouse. Vertical dashed line indicates gaze onset. A-F. Vertical scale bar: 200 °/s. Horizontal scale bar: 20 milliseconds. F. Percentage of the total eye-head co-initiated and head-initiated orientations per mouse used in the study. Totals per mouse shown. $\chi^2 = 84.09$, dof=8, $p<0.0001$. Chi-square independence test.

### Supplementary Figure 5

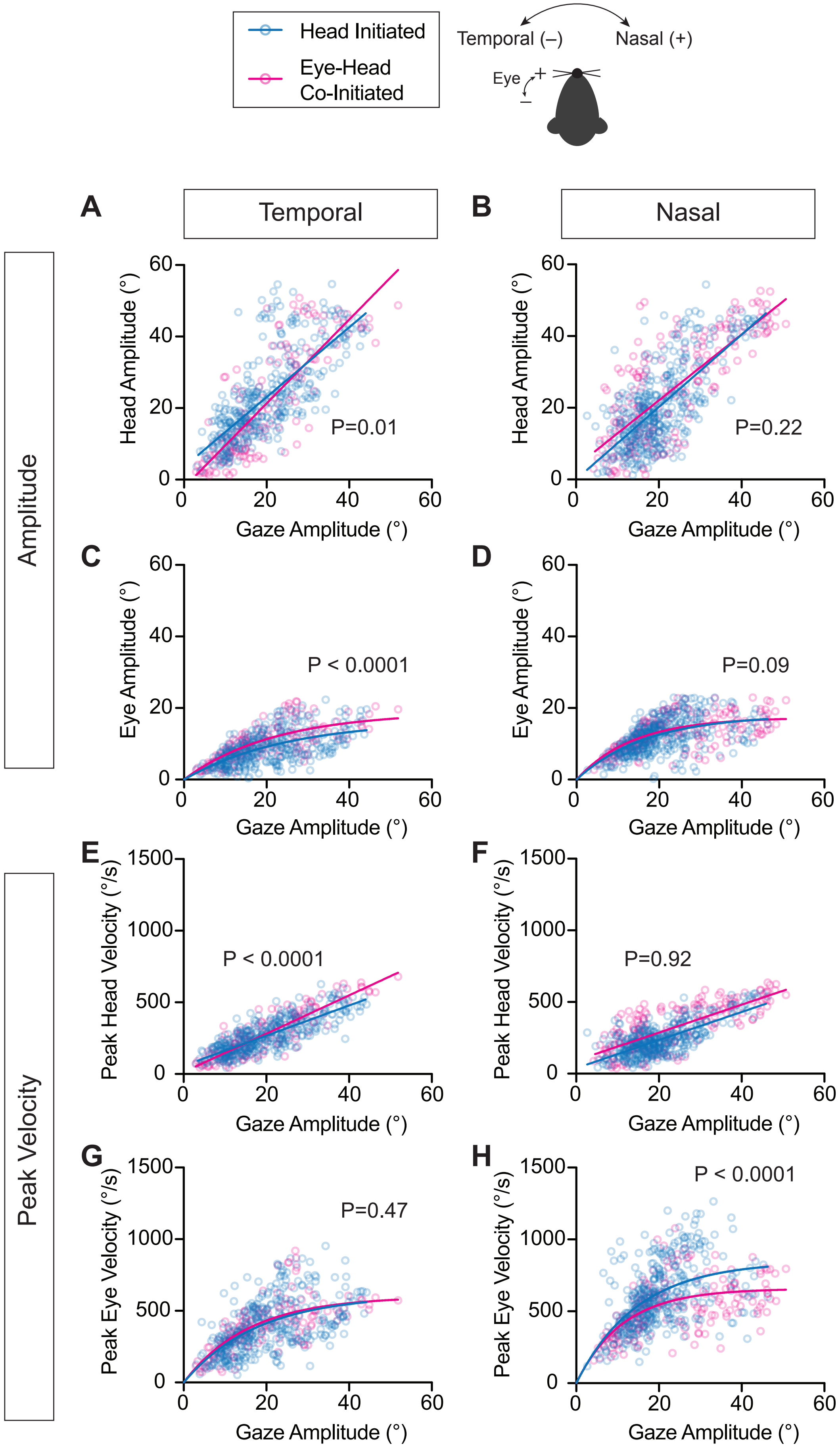

**Supplementary Figure 5: Head and eye amplitudes and peak velocities with respect to gaze amplitude, stratified by gaze redirection type. A, B.** Head movement amplitude with respect to gaze amplitude for temporally (A) and nasally (B) directed movements. Linear Regression, Temporal Head-Initiated: $R^2 = 0.56$, slope=0.98 (95% CI: [0.89, 1.06]), N = 369 segments across 9 mice; Temporal Eye-Head Co-Initiated: $R^2 = 0.68$, slope=1.17 (95% CI: [1.04, 1.30]), N = 149 segments across 9 mice; slope comparison: p=0.01. Nasal Head-Initiated: $R^2 = 0.50$, slope=1.01 (95% CI: [0.91, 1.11]), N = 391 segments across 9 mice; Nasal Eye-Head Co-Initiated: $R^2 = 0.59$, slope=0.92 (95% CI: [0.81, 1.03]), N = 194 segments across 9 mice; slope comparison: p=0.22. **C, D.** Eye movement amplitude with respect to gaze amplitude for temporally (C) and nasally (D) directed movements. Temporal Head-Initiated: $R^2 = 0.37$, $\tau$ **=** 24.9° (95% CI: [18.5, 35.7]), N = 369 segments across 9 mice. Temporal Eye-Head Co-Initiated: $R^2 = 0.57$**,** $\tau$ = 22.0° (95% CI: [16.2, 31.5]), N = 149 segments across 9 mice; curve comparison: $p < 0.0001$. Nasal Head-Initiated: $R^2 = 0.34$, $\tau$ = 16.9° (95% CI: [13.6, 21.2]), N = 391 segments across 9 mice. Nasal Eye-Head Co-Initiated: $R^2 = 0.45$, $\tau$ = 15.8° (95% CI: [13.6, 18.4]), N = 194 segments across 9 mice; curve comparison: p = 0.09**. E, F.** Peak velocity of head movement with respect to gaze amplitude for temporally (E) and nasally (F) directed movements. Temporal Head-Initiated: $R^2 = 0.66$, slope=10.5 (95% CI: [9.74, 11.3]), N = 369 segments across 9 mice; Temporal Eye-Head Co-Initiated: $R^2 = 0.79$, slope=13.4 (95% CI: [12.3, 14.5]), N = 149 segments across 9 mice; slope comparison: $p < 0.0001$. Nasal Head-Initiated: $R^2 = 0.52$, slope=9.80 (95% CI: [8.87, 10.7]), N = 391 segments across 9 mice; Nasal Eye-Head Co-Initiated: $R^2 = 0.57$, slope=9.73 (95% CI: [8.53, 10.9]), N = 194 segments across 9 mice; slope comparison: p=0.92. **G, H.** Peak velocity of eye movement with respect to gaze amplitude for temporally (G) and nasally (H) directed movements. Temporal Head-Initiated: $R^2 = 0.34$, $\tau$ = 17.7° (95% CI: [14.0, 23.1]), N = 369 segments across 9 mice. Temporal Eye-Head Co-Initiated: $R^2 = 0.53$, $\tau$ = 15.8° (95% CI: [12.8, 20.9]), N = 149 segments across 9 mice; curve comparison: p = 0.47. Nasal Head-Initiated: $R^2 = 0.28$, $\tau$ = 14.8° (95% CI: [12.0, 18.3]), N = 391 segments across 9 mice. Nasal Eye-Head Co-Initiated: $R^2 = 0.37$, $\tau$ = 11.3° (95% CI: [9.21, 13.8]), N = 194 segments across 9 mice; curve comparison: $p < 0.0001$. Blue = Head-Initiated, magenta = Eye-Head Co-Initiated.

**Supplementary Figure 6**

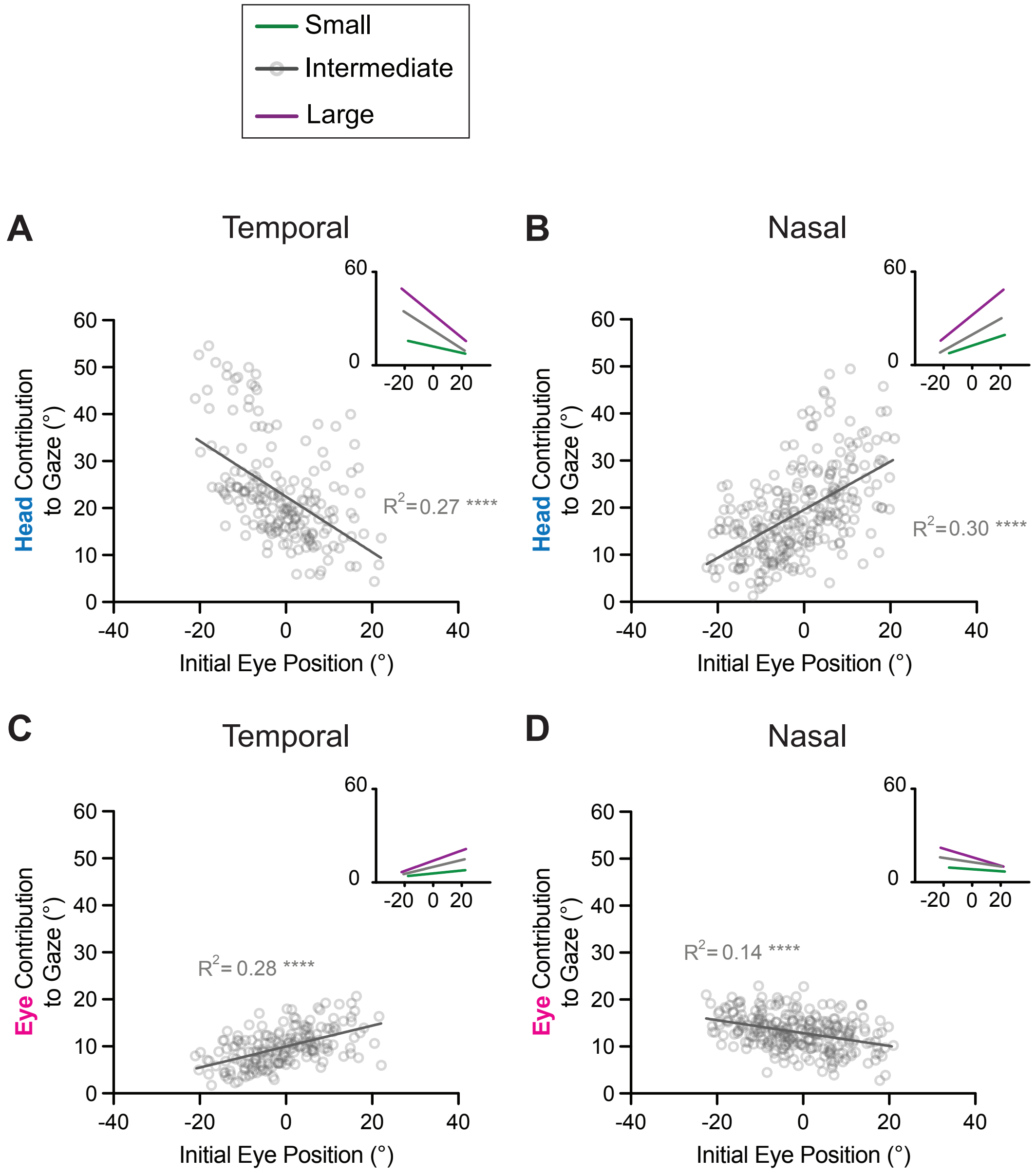


**Supplementary Figure 6: Influence of initial eye position on head and eye contributions to gaze, continued, Related to Figure 5. A, B.** Contribution of head movement to gaze with respect to initial eye position for intermediate temporally (A) and nasally (B) directed movements. Linear Regression, Head Temporal—$R^2$ = 0.27, p < 0.0001, slope = -0.59 (95% CI: [-0.73, -0.45]),N = 186 segments across 9 mice. Inset: small, intermediate, large line comparisons: F = 18.5, DFn = 2, DFd = 575, p < 0.0001; Head Nasal—$R^2$ = 0.30, p < 0.0001, slope = 0.51 (95% CI: [0.42, 0.61]), N = 267 segments across 9 mice. Inset: small, intermediate, large line comparisons: F = 14.00, DFn = 2, DFd = 634, p < 0.0001. **C, D.** Contribution of eye movement to gaze with respect to initial eye position for intermediate temporally (C) and nasally (D) directed movements. Eye Temporal—$R^2$ = 0.28, p < 0.0001, slope = 0.22 (95% CI: [0.17, 0.27]),N = 184 segments across 9 mice. Inset: small, intermediate, large line comparisons: F = 18.96, DFn = 2, DFd = 573, p < 0.0001; Eye Nasal—$R^2$ = 0.14, p < 0.0001, slope = -0.14 (95% CI: [-0.18, -0.10]),N = 265 segments across 9 mice. Inset: small, intermediate, large line comparisons: F = 17.15, DFn = 2, DFd = 632, p < 0.0001. Green = small, grey = intermediate, purple = large orienting movements. Schematic of temporal and nasal directions for head and eye shown.

**Supplementary Figure 7**

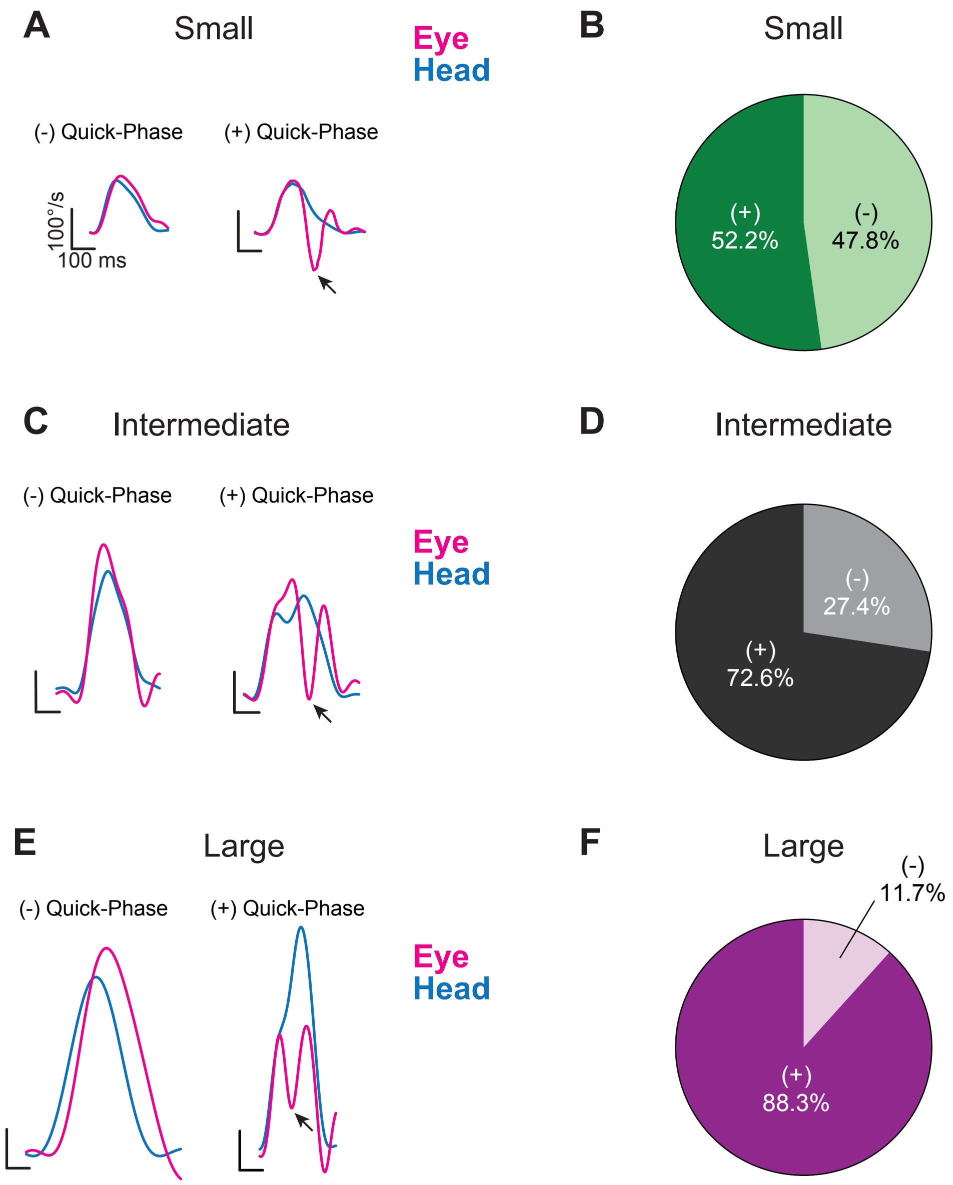


**Supplementary Figure 7: Quick-phase occurrences during passive VOR, Related to Figure 6. A, C, E.** Representative velocity traces with (+) and without (-) a quick-phase for small (A), intermediate (C), and large (E) passive gaze stabilization acquired during the passive VOR behavior paradigm. Magenta=eye (inverted for visualization). Blue=head. Arrow indicates the peak of the quick-phase. **B, D, F.** Distribution of passive gaze stabilization segments with (+) or without (-) a quick-phase. B. Small: (+) = 52.2%; (-) = 47.8%. D. Intermediate: (+) = 72.6%; (-) = 27.4%. F. Large: (+) = 88.3%; (-) = 11.7%.

### Supplementary Figure 8

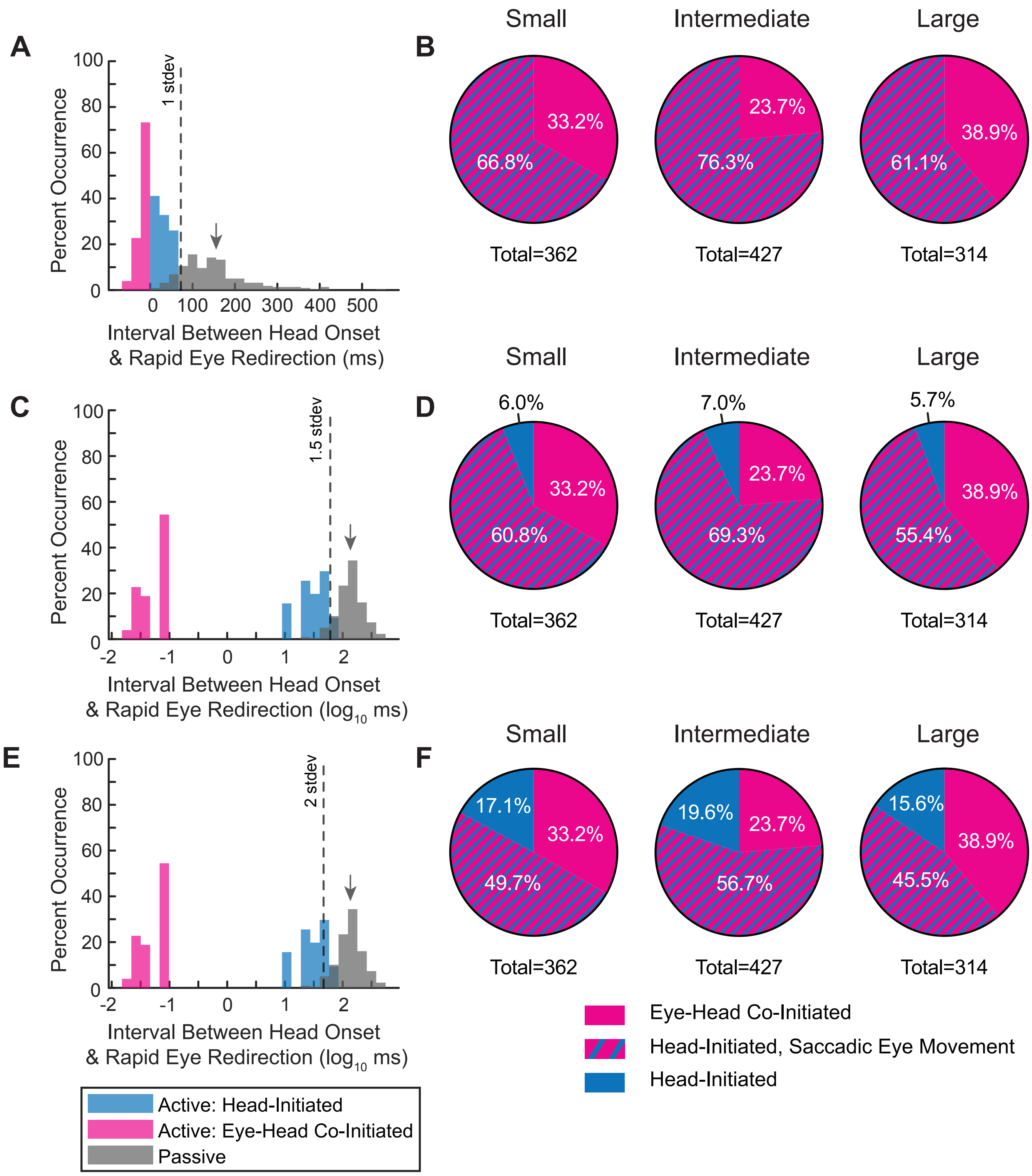

**Supplementary Figure 8: Additional cutoff metrics for comparison of active: head-initiated and passive with quick-phase, Related to Figure 6. A.** Same as Figure 6C (Active: Eye-Head Co-Initiated, Active: Head-initiated and Passive with Quick-Phase). Dashed line indicates 1 standard deviation away from the mean of the passive with quick-phase. 100% of the active: head-initiated data falls outside of 1 standard deviation of the mean of passive with quick-phase. **B.** Replotting of head-initiated versus eye-head co-initiated distributions shown in Figure 2 G-I, now identifying the subset of head-initiated gaze shifts demonstrating saccadic eye movement with a 100% cutoff. Small: 33.2% eye-head co-initiated, 66.0% head-initiated with saccadic eye movement, 0.0% remaining head-initiated. Intermediate: 23.7% eye-head co-initiated, 76.3% head-initiated with saccadic eye movement, 0.0% remaining head-initiated. Large: 38.9% eye-head co-initiated, 61.1% head-initiated with saccadic eye movement, 0.0% remaining head-initiated. **C.** Same data as (A), now log transformed. Dashed line indicates 1.5 standard deviations away from the mean of passive with quick-phase. 90.6% of the active: head-initiated data falls outside of 1.5 standard deviations of the mean of passive with quick-phase. **D.** Replotting of head-initiated versus eye-head co-initiated distributions, now identifying the subset of head-initiated gaze shifts demonstrating saccadic eye movement with a 90.6% cutoff. Small: 33.2% eye-head co-initiated, 60.8% head-initiated with saccadic eye movement, 6.0% remaining head-initiated. Intermediate: 23.7% eye-head co-initiated, 69.3% head-initiated with saccadic eye movement, 7.0% remaining head-initiated. Large: 38.9% eye-head co-initiated, 55.4% head-initiated with saccadic eye movement, 5.7% remaining head-initiated. **E.** Same as in (C), dashed line indicates 2 standard deviations away from the mean of passive with quick-phase. 74.0% of the active: head-initiated data falls outside of 2 standard deviations of the mean of passive with quick-phase. **F.** Replotting of head-initiated versus eye-head co-initiated distributions, now identifying the subset of head-initiated gaze shifts demonstrating saccadic eye movement with a 74.0% cutoff. Small: 33.2% eye-head co-initiated, 49.7% head-initiated with saccadic eye movement, 17.1% remaining head-initiated. Intermediate: 23.7% eye-head co-initiated, 56.7% head-initiated with saccadic eye movement, 19.6% remaining head-initiated. Large: 38.9% eye-head co-initiated, 45.5% head-initiated with saccadic eye movement, 15.6% remaining head-initiated. Arrows in A, C, and E indicate mean of passive with quick-phase. Histogram bin size = 22.2 milliseconds, corresponding to two video frames.
